# Make like a tree and leave: How will tree species loss and climate change alter future temperate broadleaved forests?

**DOI:** 10.1101/2023.06.26.546609

**Authors:** Bede West, Davey L. Jones, Emma L. Robinson, Aidan M. Keith, Simon Kallow, Robert H. Marrs, Simon M. Smart

## Abstract

Forest creation has the potential to reduce biodiversity loss and mitigate climate change but, tree disease emergence may counteract this. Further, given decadal timescales required for forest establishment, climate change is increasingly likely to act as a filter on plant community assembly. In the temperate lowlands succession takes 30 to 50 years for non-forest land to establish woodland plant assemblages, while the timescales required for new forest to sequester carbon suggest unassisted succession will be too slow for net zero 2050 targets. However, if plantations can establish faster than succession it would be beneficial to recommend planting native species as soon as possible. We explore scenarios of broadleaved woodland development across Wales, UK, as a case study area. We use a suite of empirical species niche models for British plants to estimate the potential species composition of forests with, and without, projected climate change. Additionally, we examine how tree canopy composition alters if *Fraxinus excelsior* is widely impacted by ash-dieback (*Hymenoscyphus fraxineus*). The results suggest soil total carbon and nitrogen could achieve baseline broadleaved forest values in less than 30 years. However only timber and woody flora species groups showed diversity surpassing baseline broadleaved forest diversity, with nectar plants and ancient woodland indicator species failing to reach baseline equivalents within 30 years; although complete congruence is unlikely given baseline forests could be hundreds of years old. Where *Fraxinus excelsior* was removed from the species pool we predicted that a scrub phase will persist or, if present, *Acer pseudoplatanus* will become the canopy dominant. The heavier shade cast this species is likely to result in differences in species composition of the understory and ground flora diversity is likely to decrease. Reliance on unassisted succession will also depend critically on (a) dispersal from local source populations and (b) on establishment filters that could be severe in landscapes with high management intensity history. These findings indicate that leaving the UK’s fragmented habitats to relying on already degraded successional processes could lead to poor afforestation outcomes.

**Highlights:**

- Afforestation can mitigate global change but tree disease makes outcomes uncertain
- Afforestation methods establishment timescales and time for benefits to occur
- We model afforestation and predict how soils and plants change with climate
- Ash loss from die-back is replaced by low low-canopy woodland / scrub over 30 years
- Afforestation achieves baseline forest values for some variables within 30 years

## 1 Introduction

Afforestation is considered a major part of global attempts to sequester carbon and help each country achieve net zero (Read et al., 2009; Di Sacco A & Hardwick K et al., 2020; Stafford et al., 2021; HM Government, 2021).) Given the long-term nature of forest establishment, there is an obvious need for information about the likely results that will occur under a range of future scenarios. Such scenarios will include projected climate change, and the potential for increased death of dominant tree species through the advent of new diseases (Mitchell et al., 2016, Skovsgaard et al., 2017). The obvious way to obtain such predictions is through modelling the responses of forest species. This approach must be able to predict likely colonisation of species which are not necessarily present in the forest at the start but are available in the local species pool. This pool of potential colonisers is dark diversity (Partel, et al., 2011).

Here, we introduce a modelling approach that takes into account:

A. Future climate change scenarios, as this is increasingly likely to become a novel filter on species composition of the canopy and understorey potentially giving rise to vacant niche space or no-analogue assemblages (Read et al., 2009; Alexander et al., 2016).
B. Reduced acid deposition, as soil pH is recovering from atmospheric deposition (Emmett et al., 2010)
C. Death of a dominant tree species, likely to cause change to forest habitats (Mitchell et al., 2016)
D. The potential for colonisation from new species from outside the starting plant community (the dark diversity; Pärtel et al., 2011) and the time this takes to reflect baseline forest conditions (20-50 years; Walker, Sparks and Swetnam, 2000; Harmer et al., 2001; Poulton et al., 2003; Ashwood et al., 2019)

Ash (*Fraxinus excelsior* L.) is the tree we remove from the species pool due to mortality from ash-dieback (*Hymenoscyphus fraxineus* (T.Kowalski) Baral, Queloz & Hosoya), which has been in western Europe for around a decade (Kjær et al., 2012; Pautasso et al., 2013; Baral, Queloz and Hosoya, 2014). We produce predictions for forests in the Environment & Rural Affairs Monitoring and Modelling Programme (ERAMMP, Emmett *et al*., 2016) for the Glastir agri-environment scheme in Wales (Rose, 2011; Welsh Government, 2017). Allowing us to make a country-wide predictions. Our models produce estimates on carbon sequestration and potential biodiversity change.

### 1.2. Forest carbon sequestration

As afforestation sequesters carbon in both above (Di Sacco, & Hardwick, et al., 2020) and below ground pools (Minasny et al., 2017; Mayer et al., 2020), whilst also being profitable and mitigating biodiversity loss (Di Sacco, & Hardwick, et al., 2020; Read et al., 2009). Speeding up afforestation, therefore, seems desirable. However, Read et al. (2009) highlighted, over 10 years ago, that benefits from UK forest establishment may take 50 to 100 years to take effect; too slow for UK 2050 targets (HM Government, 2021).

Two key factors have been identified as important to ensure net C gain occurs in forests. The first considers soil C content and where not to plant. Planted forests on high C soils often fail to show a net C gain even decades after planting, and sometimes suffer C loss (Friggens et al., 2020; Mayer et al., 2020; Casado et al., 2022). Planting on these soils must be avoided, this excludes acid-grasslands and heathland habitats from planting in the study area (Emmett et al., 2017 & Seaton et al., 2020; *SM 1* for Welsh broad habitat details). The second, is C priming where C introduced into the soil under increased CO_2_ and higher temperatures causes greater microbial activity, respiration and hence loss of soil C (Smith et al., 2013). Any increase in soil C especially in the upper layers (e.g. top 15 cm considered here) may not be representative of an overall increase in soil C sequestration, certainly if soil fertility is low (De Graff et al., 2006; Hungate et al., 2009). This is because labile C can prime microbial activity in lower layers giving rise to an overall C loss (Smith et al., 2013; Guenet et al., 2018). However at multi-year timescales increases in the C pool provide net C sequestration, at > 3 years this tends to be increases in the labile fraction (De Graff et al., 2006; Guenet et al., 2018; Mayer et al., 2020). Thus, for more recalcitrant C pool gains > 25 years is more realistic (Cotrufo et al., 2013).

### 1.3. Biodiversity and carbon

As benefits to biodiversity and C sequestration may not be achievable in the same area, strategies are required to optimise management interventions, taking account of the starting landscape, and trade-offs within ecosystem and land use types (Read et al., 2009; Di Sacco et al., 2020; Linney et al., 2020). We consider forest establishment by progress towards established (baseline) forest conditions, via modelled changes in plant communities and soil variables (pH, carbon, and nitrogen) which have been highlighted as important for soil health in Seaton et al. (2020). We use a 30-50 year UK forest establishment time-frame, which is longer than the 20-30 years consider to be a possible establishment time elsewhere (Falkengren-Grerup, et al., 2006; Vesterdal et al., 2008; Brunet et al., 2012; Thomaes et al., 2012). However, there is evidence that many forest species can establish within 15 years when adjacent to older forest (Brunet et al., 2012). If faster forest establishment is achievable, then faster benefits will occur.

One major constraint to the assembly of woodland plant communities is the depletion of plant species pools due to the degradation of UK landscapes with only 13% forest cover, one of the lowest in Europe (Hayhow et al., 2019; Forest Research, 2020). Natural migration of understory forest species may occur at <3 to 12 m per year depending on species (Brunet et al., 2012). But tree colonisation in the Welsh uplands in Wales (McGovern et al., 2013) and northern-England (Marrs et al. 2018) is slow with no tree cover established after 50-60 years in grazing exclusion experiments. Even when wider upland species diversity has been seen to increase in that time (Alday et al., 2022) it is clear that the way a forest is established, and the age it is allowed to reach before it is altered is a major factor in its, C sequestration, biodiversity, and therefore health (Brunet et al., 2012; Kröel-Dulay et al., 2015; Ashwood et al., 2019; Berdeni, Williams and Dowers, 2021). If plantation broadleaved forest can establish faster than successional timeframes (< 30 years), it appears best to recommend planting native species using minimum soil disturbance techniques (*sensu* Berdeni et al., 2021) to mitigate climate change and biodiversity loss.

### 1.4. Research aims

We explored the combined effects of climate change and disease-induced tree mortality by modelling soil and climate changes (Fig. 1). This generated habitat suitability scores for woodland plants across a range of habitats in Wales, we summed these at high-resolution to estimate dark diversity (Pärtel et al., 2011). Dark diversity refers to the species within the local pool that could inhabit a patch given its observed or modelled conditions (Pärtel, Szava-Kovats and Zobel, 2011). This means we model potential colonising species into future scenarios that are both observed and unobserved at baseline to provide dark diversity predictions. We applied “worst-case-scenario” (RCP 8.5) climate projections (Lowe et al., 2018; Met Office Hadley Centre, 2018) and model over longer time-periods than achieved to date (Mitchell et al., 2016).

**Fig.1.**
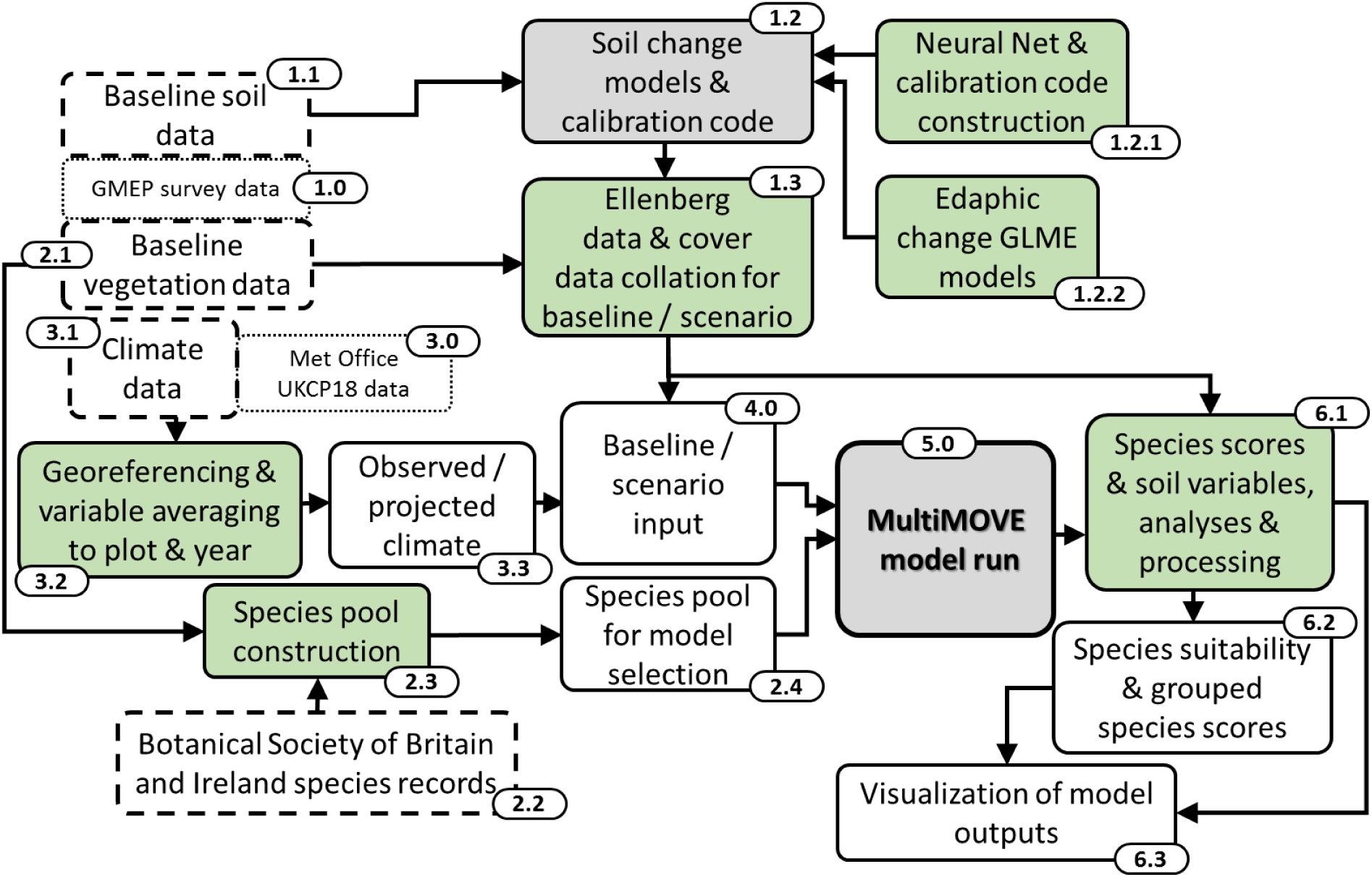
Schematic representation of the modelling workflow. Numbered boxes cross-reference to main text as numbers coded in [blue]. Green boxes represent a coded process; grey boxes represent model runs; white boxes are datasets; white boxes with dashed outlines represent input data.

Thus, we aim to produce predictions that will guide ecological restoration accounting for place-based pools of possible colonists (Pärtel et al., 2011) including: ancient woodland indicator (AWI) species (Glaves et al., 2009); woody flora (Kallow, 2014; Trivedi and Kallow, 2017); timber species (Pyatt et al., 2001; Bathgate et al., 2011); and nectar producing species (Smart et al., 2017; Alison et al., 2021). We addressed the following research questions:

i. Do modelled changes in soil conditions and increasing shade from a growing tree canopy filter forest plant assemblages over a 30-year period approximate to reference baseline conditions in established broadleaved forest?
ii. Which trees and shrubs could replace F. excelsior with and without alteration of climate?
iii. To what extent are these modelled assemblages changed in their species composition by projected climate change?

## 2. Methods

The following sections follow the numbering in Fig. 1, detailed in the parentheses. All modelling, statistical analysis and data plotting was conducted in R statistical software version 4.0.3 (R Core Team, 2019). The R package MutliMOVE (Fig. 1: 5.0) forms the ecological niche modelling core of the workflow (Henrys et al., 2015).

### 2.1. GMEP Survey data [1.0]

Co-located soil and plant species compositional data were recorded as part of the Glastir Monitoring and Evaluation Program (GMEP); 732 plots (200 m^2^) surveyed once between 2013 and 2016 were used; detailed methods in Emmett et al. (2017) and Seaton et al. (2020).

### 2.2. Baseline soil sampling [1.1]

Five soil cores (5 cm diameter x 15 cm depth) were taken from the edge of the central 2×2 m within each plot. Soil pH in water, carbon (%), nitrogen (%) and gravimetric moisture content (%) from each plot define baseline model inputs and are the starting values that are adjusted or modelled over-time, representing tree planting and subsequent woodland development.

### 2.3. Soil response models and calibration [1.2]

A suite of transfer functions were used to estimate values of vegetation indicator variables (mean Ellenberg values) for each quadrat given measured soil variables (Ellenberg et al. 1992). This step was necessary because the plant ecological niche models (ENM) were originally built using mean Ellenberg values plus vegetation height and climate as inputs (De Vries et al., 2010; Smart et al., 2010; Henrys et al., 2015). Ellenberg values represent the position of a plant species along ecological gradients of wetness (Ellenberg F), fertility (Ellenberg N) and reactivity (Ellenberg R) created by Ellenberg et al. (1992) and modified for Britain by Hill et al. (2004). To avoid circularity in building plant species niche models we excluded the focal species from the calculation of mean Ellenberg scores when modelling its niche (*see*, SM 2.4).

#### 2.3.1. Neural Net calibration code [1.2.1]

Neural networks were used to model the three Ellenberg scores using soil variables in 2.1.2 as inputs (*sensu,* West et al. 2023). Details of the Neural networks are presented in SM 2.4.

#### 2.3.2. Edaphic change models [1.2.2]

Our approach to modelling change in the soil variables over time in response to planting and woodland development was data driven. Soil change under woodland development was modelled as follows; a literature search was conducted to find empirical soils data from time-series or chronosequences that measured the impact of afforestation on soils by UK native broadleaved species. Care was taken to ensure soil variables were measured in the same way as GMEP soil measurements. This yielded nine data sets which were either sourced from the published literature or provided by the original authors (see, SM 2, Table SM.1). These generalised linear mixed effect (GLME) models were based on the following covariates: starting value of the variable modelled, time (years of woodland development) and afforestation type. All models used a random effect for study or data source. GLME models differed in terms of the final set of covariates included. GLME construction and training data are presented in SM 2.

An annual incremental addition to soil pH was added following West et al. (2023) and Emmett et al. (2010) for each broad habitat type to represent a recovery from deposition-based acidification, reflecting the expected response to ongoing recovery from historically-high levels of sulphur deposition (Kirk et al., 2010).

### 2.4. Ellenberg and cover data collation [1.3]

Soil measurements for each GMEP plot were inputted to the neutral network models and translated into predicted mean Ellenberg values, these are the required inputs for the MultiMOVE ecological niche models (ENM). Baseline cover-weighted canopy height was calculated using vegetation species data (2.1) and average vegetative height (Hill et al., 2004; West et al. 2023). This derived variable expresses the successional stage of the habitat and, by proxy, the light availability at plot ground level (Depauw et al., 2020).

To represent forest growth, we incrementally increased cover-weighted canopy height (CWCH) by the years modelled in the workflow (Table 1, B). A literature review was conducted to find growth rates for *F. excelsior* and other broadleaves grown in and native to the UK. Where *F. excelsior* was absent, an average across the native broadleaved species present was taken (see the following for data: Claessens et al., 1999; Hein, 2003; Harmer et al., 2005; Dobrowolska et al., 2011; Harmer et al., 2012). The CWCH year increments (Table 1) were created by taking an average height (m) reached over the given number of years from the literature categorised as CWCH values. The categorisation of meters to CWCH was as for the MultiMOVE manual, i.e., CWCH 5 = 1.0-3.0 m; CWCH 6 = 3.1-6.0 m; CWCH 7 = 6.1-15.0m; CWCH 8 >15m (Hernys et al., 2015). Half units were used in 2026 & 2036 as growth rates were not always sufficiently different to move between categories, but growth still had to be represented. Model testing showed that this approach to filtering species by their canopy height and by soil conditions was able to reproduce the observed woodland community type satisfactorily, given succession (see Fig. 11.4, Rowe et al., 2015)).

**Table 1.** Details of woodland development scenarios modelled with and without *Fraxinus excelsior*. Part A of the table refers to specifics of the scenarios, part B shows the years within the scenarios. Scenarios are: *Baseline* = Modelled forests from observed environmental data in GMEP survey; *Baseline CC* = as for Baseline but with the predicted climate change data as inputs; *Plantation broadleaved* = Broad habitat types, Arable and Horticulture, Improved Grassland, Neutral grassland and Bracken with soils modelled as having been planted with broadleaved trees under a baseline climate; *Plantation broadleaved CC* = As for *Plantation broadleaved* but with the predicted climate change data as inputs. Baseline climate = Baseline average climate (1981 to 2016); Predicted climate = High emissions.

| Scenario | Management | Species pool | Soil models categorical setting | Recovery from acidification applied | Scenario versions. [years modelled] |
| --- | --- | --- | --- | --- | --- |
| *Baseline* | Observed forests in GMEP survey. | Dark diversity; No *F. excelsior* | None | No | 2, Observed climate averaged (1981 to 2016) with/without *F. excelsior*. [2016] |
| *Baseline CC* | Observed forests in GMEP survey. | Dark diversity; No *F. excelsior* | None | Yes | 18, Predicted climate, with/without *F. excelsior*. [2021,2026,2029,2036,2043,2046,2050,2056] |
| *Plantation broadleaved* | Planting UK native broadleaves | Dark diversity; No *F. excelsior* | *pH & N%*: planted broadleaved<br>*C%*: broadleaved | Yes | 18, Observed climate, with/without *F. excelsior*. [2021,2026,2029,2036,2043,2046,2050,2056] |
| *Plantation broadleaved CC* | Planting UK native broadleaves | Dark diversity; No *F. excelsior* | *pH & N%*: planted broadleaved<br>*C%*: broadleaved | Yes | 18, Predicted climate, with/without *F. excelsior*. [2021,2026,2029,2036,2043,2046,2050,2056] |

B
| Years from Baseline (2016) | Time interval description | Cover weighted canopy height setting | Year modelled |
| --- | --- | --- | --- |
| 5 | Post planting stage | 5 | 2021 |
| 10 | First decade | 6.5 | 2026 |
| 13 | Longest UK agri-environment funding time (Rose, 2011) | 7 | 2029 |
| 20 | Second decade of change | 7.5 | 2036 |
| 27 | The 25-year environment plan end | 8 | 2043 |
| 30 | Third decade | 8 | 2046 |
| 34 | Net zero emissions goal year | 8 | 2050 |
| 40 | Fourth decade | 8 | 2056 |

### 2.5. Baseline vegetation [2.1]

Baseline vegetation data was equivalent to that used in West et al., (2023), species and their percentage cover for broad habitats: Arable and Horticulture (A&H); Improved Grassland (IG); Neutral Grassland (NG); and Bracken (Br). *See, SM 1,* for broad habitat type descriptions.

### 2.6. Defining the wider species pool [2.2, 2.3]

The Botanical Society of Britain and Ireland distribution database provided the local 10 km species pool (BSBI, 2018) as defined in West et al., (2023). The species pool modelled at each plot location comprised the unique list of plant species observed in the baseline quadrats (Fig, 1.0 & 2.1), those recorded in other quadrats in the GMEP 1 km square (Fig 1, 2.1) and additional species present in the wider 10 km square pool.

### 2.7. Observed and predicted climate data [3.0, 3.1]

Here, we used the same climate data as West et al., (2023) i.e., observed historic data averaged from 1981 to 2016 as a baseline; and ‘worst case scenario’ future high emissions (2021-2056; RCP8.5), both at the 1 km cell resolution (Lowe et al., 2018; Met Office et al., 2019; Robinson et al., 2022). The RCP8.5 1 km data source is described in *SM 4*.

### 2.8. Georeferencing and variable averaging [3.0, 3.1]

Climate data were geo-referenced to each plot location via the 1 km climate data cell they occurred within, see *SM 4*.

### 2.9. Observed or projected climate application [3.3]

The climate data was used in the three scenarios, with all modelling done for 2016 using baseline climate data, the planted scenarios applied baseline and projected climate so that results for each, across years (Table 1), could be compared.

### 2.10 Model inputs [4.0]

Lastly, the MultiMOVE model was run for all species in the pool attached to each quadrat location using the seven input variables (three mean Ellenberg values; cover weighted canopy height; minimum January and maximum July temperature, and total annual precipitation). MultiMOVE then uses these inputs to predict habitat suitability scores for the species in each modelled quadrat location. These habitat suitability scores were then ranked, providing a prediction of which species will be filtered in (or out) of plots under the differing scenario inputs (Table 1, A).

### 2.11. MultiMOVE [5.0]

The MultiMOVE ENM consists of an ensemble of five modelling methods whose outputs are combined to produce a weighted model average habitat suitability score. The ensemble models the realised niche of 1262 taxa within Great Britain covering the most common and many less common plants and bryophytes (see Henrys et al., 2015; Smart et al., 2019 for full description). The ENM package has been utilised by many studies under multiple scenarios and is subject to a process of ongoing validation and testing (West et al., 2023; De Vries et al., 2010; Henrys, Smart, et al., 2015; Rowe et al., 2015; Emmett et al., 2017; Smart et al., 2019).

### 2.12. Summarising model outputs [6.1]

Outputs from Fig. 1, 1.3 & 5.0 were summarised by habitat type and scenario for the modelled interval 2016 to 2056. We used an unpaired t-test on loge-transformed data to compare soil variables against the baseline broadleaved woodland values This allowed determination of how modelled planted soil for a given year differed from observed forest soils.

We then used logistic regression to test whether species’ presence/absence in the baseline was correlated positively with the modelled habitat suitability for each plot location (*see, SM 3*).

### 2.13. Habitat suitabilities and species groups [6.2]

Habitat suitability scores were analysed in two ways. First, we summed suitability scores for specific subsets of ecosystem service-supporting species yielding estimates of dark diversity (see, Fig. 2) for each quadrat location, year, habitat and species group (Calabrese et al., 2014). Dark diversity (Pärtel, Szava-Kovats and Zobel, 2011) refers to all the species within a local area of a site that could grow under the environmental conditions at the site, here this includes species pools in Fig. 1, 2.1 & 2.2. Species with higher habitat suitability scores are, therefore, those best suited to soil, climate and vegetation height at each location. We produced dark diversity estimates for four species groups: the UK woody flora, comprising natives trees and shrubs (Kallow, 2014; Trivedi and Kallow, 2017); ancient woodland indicators for Wales (Glaves et al., 2009); commercially-valuable broadleaved timber species, i.e., a timber-producing subset of the woody flora (Pyatt, Ray and Fletcher, 2001; Bathgate et al., 2011); and nectar producing species (Smart et al., 2017; Alison et al., 2021). Species group lists are within*, SM 3.2, Table SM.3*. Second, the habitat suitability scores per plant species were summarised as a frequency table enabling matching to the British National Vegetation Community (NVC) types (Rodwell, 1998) via the software MAVIS (Smart, 2000; West et al. (2023).

**Fig.2.**
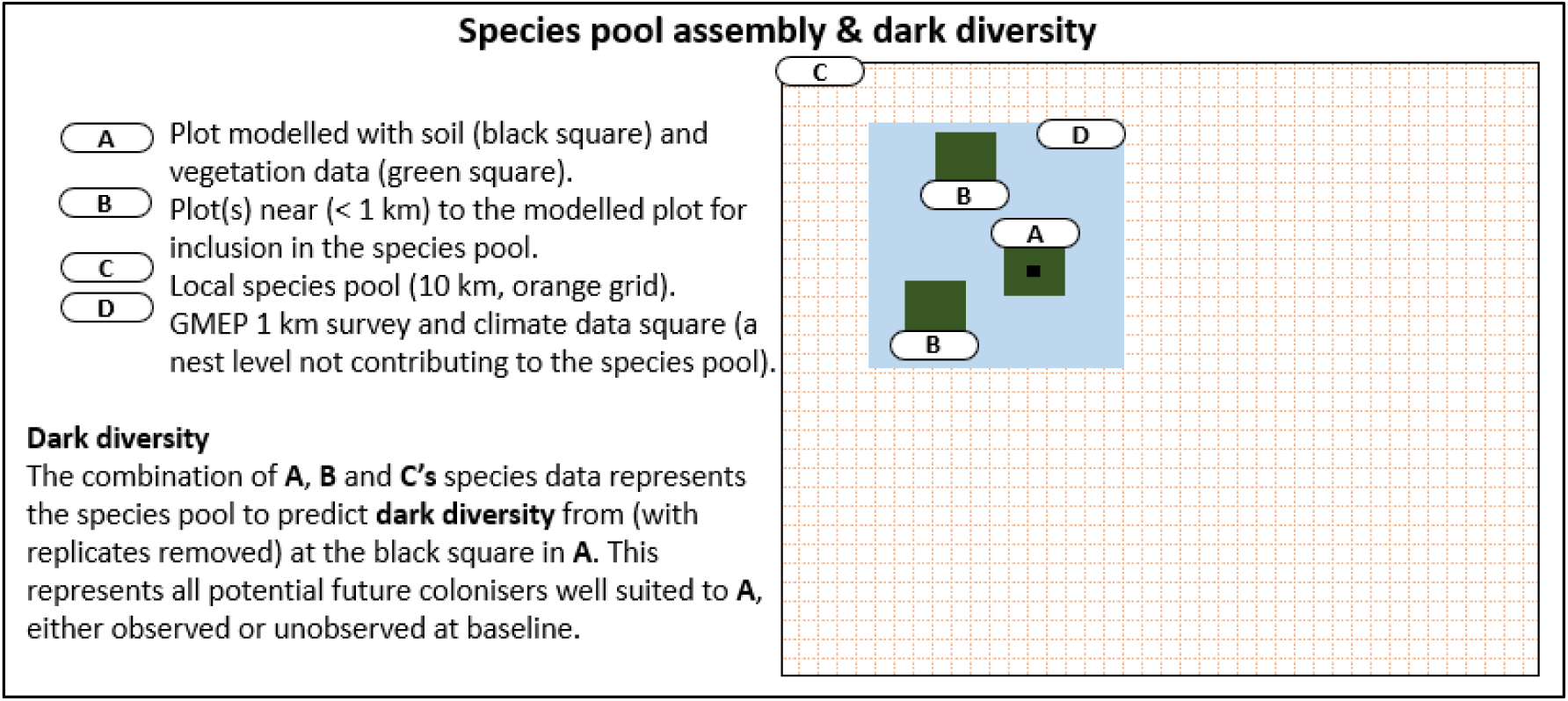
Assembly of the species pool contributing to dark diversity. The spatial nesting (not scaled) of the squares provides a visual as well as descriptive explanation of dark diversity. The survey square within C refers to the Glastir Monitoring and Evaluation Program 1 km squares representing land use across Wales.

A key component of the workflow for considering *H. fraxinus* impacts is the inclusion or removal of *F. excelsior* from the model outputs. Removal of *F. excelsior* from the outputs allows for species and vegetation types ranked below *F. excelsior* vegetation to be identified as likely replacements and, hence, how this may change the future vegetation.

### 3.0. Results

### 3.1. Soil variables modelled

Outputs of the soil variables models (Fig. 3), show all habitats are expected to increase in pH given alterations applied due to expected recovery from historically high S deposition. Moreover, every habitat (Fig. 3) other than bracken starts at higher pH than the broadleaved woodland reference (Fig. 3, A). Nitrogen and carbon in some planted broad habitats appear to respond with values becoming congruent with, or not significantly different from, the reference baseline broadleaved habitat by the 2040s.

**Fig. 3.**
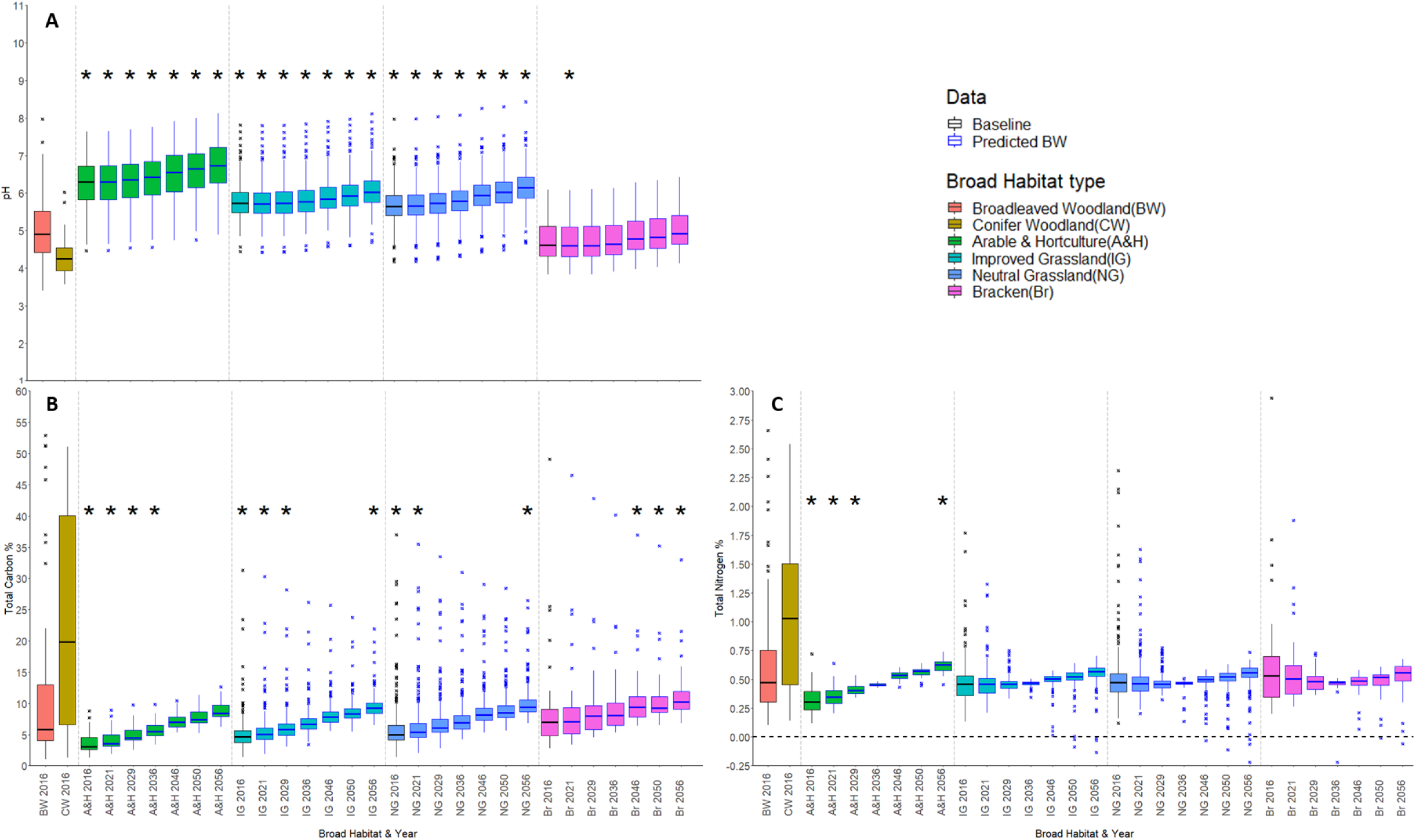
Modelled change in soil variables under tree planting per Broad Habitat (BH) type and year, for pH (A), total soil carbon percentage (B) & total soil nitrogen percentage (C). The year 2016 for each BH type represents the observed data (black boxes), subsequent years data were predicted by using 2016 data as inputs to generalised linear mixed effect models of broadleaved woodland plantation on respective soil variables (blue boxes). Asterisks (*) at Y=9 for pH; Y=40 for Carbon & Y=2 for Nitrogen, represent significant differences (p-value=0.05) of logged variables of the 2016 Broadleaved Woodland baseline compared to the modelled values for the Broad Habitat type and year below each asterisk. Where asterisks are not shown planted habitat values were not significantly different from the target Broadleaved Woodland baseline.

Planting is predicted to increase topsoil C concentrations (Fig. 3, A), habitats either exceed or reach equivalent baseline broadleaved reference values by the mid-2030s. Bracken is similar to the baseline at the start but exceeds broadleaved woodland values by 2046. Arable, improved and neutral grassland start with significantly lower values at baseline but are then predicted to accumulate carbon becoming no different (arable) or exceeding (grasslands) the baseline by 2056.

Little change in soil N is predicted, other than in arable where values start lower than the broadleaved reference and end significantly greater (Fig. 3). Overall, the results predict increasing soil pH and C:N ratio. The higher pH starting points consistent with establishing woodland on fertile, agriculturally-managed grasslands, suggest that developing woodland plant communities will reflect higher pH than the average Welsh woodland (Fig. 3, A).

### 3.2. Model validation against baseline observations

Logistic regression showed that a greater habitat suitability score (weighted model average) increased the probability of the species being observed in baseline quadrats (*P* <0.001, *see, SM 3, Fig. SM.2*).

### 3.3. Species groups and predicted dark diversity

Predicted dark diversity for two groups of ecosystem service-supporting plants both showed deviations from the baseline broadleaved woodland across the time interval.

#### 3.3.1. Nectar plants and ancient woodland indicators

When climate change is included as a filter on the species pool, predicted diversity of both groups matches the broadleaved baseline by 2050 for Neutral Grassland and bracken starting points (Fig. 4). This also occurs for nectar plant diversity in the two agriculturally-intensive habitats, arable & horticulture and improved grassland, but woodland specialist diversity still lags behind the broadleaved reference for these two habitats with climate change included (Fig. 4). Without climate change, predicted diversity shows a persistent lag behind the baseline reference even by 2050 for both plant groups. This applies across all starting habitats except neutral grassland where nectar plant diversity moves appreciably toward, but does not match, baseline values by 2050.

**Fig. 4.**
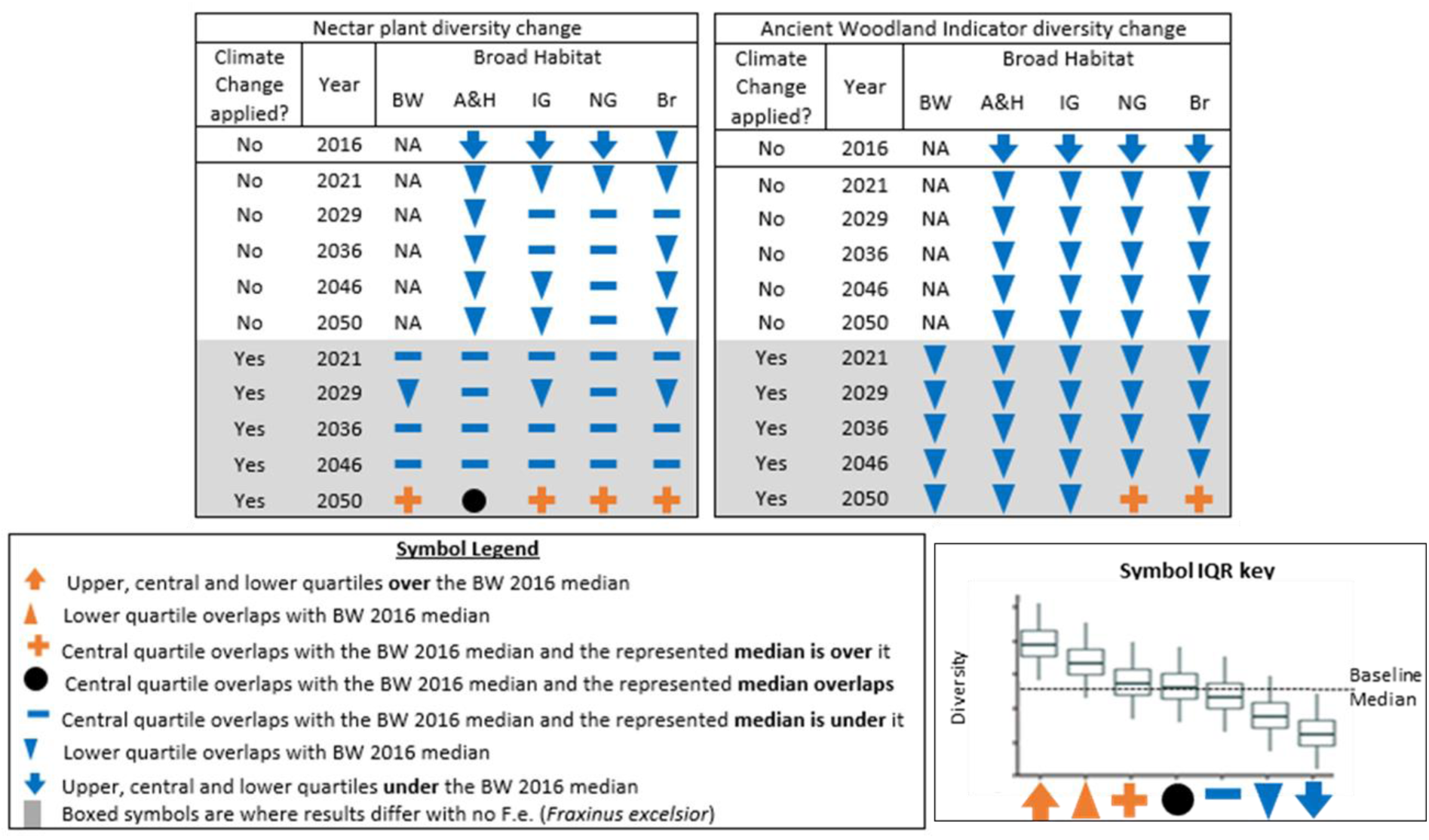
Species group diversity change. Symbols define how the species group’s diversity score (interquartile range (IQR) and median) for a given year and broad habitat type compares with the Broadleaved Woodland baseline median (2016) given accumulation of woodland dark diversity, either with or without climate change. Thus, relative to the baseline, blue symbols indicate a decrease, black, no change and orange an increase. Broad Habitat types: BW = Broadleaved woodland (2016 baseline); A&H = Arable and horticulture; IG = Improved grassland; NG = Neutral grassland; Br = Bracken. The data used to create these boxplots was generated using an ecological niche model MultiMOVE, inputs were altered to represent baseline (1981-2016) and future climates using downscale UKCP18 climate data, incremental increase of cover-weighted canopy height representing tree growth and generalised linear mixed effect models of soil variable change under broadleaved plantation. All diversity scores are representative of modelled dark diversity, thus the baseline score will be higher than the 2016 observed diversity. *See, SM 3.2, Fig. SM.8* for boxplots showing the IQRs of each species groups diversity scores per habitat and scenario.

Nectar-producing plant diversity in broadleaved woodland drops below the baseline median in Fig. 4, from 2021 to 2046; this appears to be a reshuffling of the community rather than a multi-taxa climate response. This is reflected in habitat suitability of *Hedera helix* and *Rubus fruticosus* (*SM 3.2, Fig. SM.5, B&C*) in 2021, 2036 and 2046 being below the 2016 baseline; and *Crataegus monogyna* (*SM 3.2, Fig. SM.5 A*) and *Hyacinthoides non-scripta* (*SM 3.2, Fig. SM.4, C*) rising above the baseline until 2029 after which their scores vary around the baseline until 2050 where they are higher. Under climate change, ancient woodland indicator species (AWI) in baseline broadleaved woodland showed a drop in diversity, alongside a drop in all the climate variable values in 2029 (Fig. 4 & *SM 3.2, Fig. SM.11*). This climate response is confirmed by habitat suitability of *Arum maculatum, Circaea lutetiana*, *Hyacinthoides non-scripta* and *Oxalis acetosella* all having their lowest, or near lowest, scores in 2029, then rising from 2040 onwards (*SM 3.2, Fig. SM.4*). However, no modelled planted habitat reaches the AWI median prior to 2050.

#### 3.3.2. Timber Species and Woody Flora

Dark diversity scores of both groups in Fig. 4 are predicted to have exceeded the baseline reference by 2050 with, and without, climate change. Different levels of response were observed among habitats: Bracken was predicted to show the largest positive response over the shortest period. Increasing canopy height in this habitat results in dark diversity of both species groups exceeding the modelled baseline by 2029 (Fig. 4). In broadleaved woodland filtered by climate change, similar suitability changes occur for the predicted dominants. Thus, *Acer pseudoplatanus*, *Fagus sylvatica*, *Fraxinus excelsior* and *Quercus* spp. all increase in suitability from year to year (*SM 3.2, Fig. SM.6*).

For both planting scenarios, under baseline and predicted climate (Fig. 5), there is less variation in the woody flora (*SM 3.2, Fig. SM.10,C&D*) than the subset group of timber species (*SM 3.2, Fig. SM.9, C&D*). However, individual species under climate change showed changes in the rank order of their habitat suitabilities. For example, *Corylus avellana* showed its lowest scores in 2029 & 2050 (*SM 3.2, Fig. SM.7*); but *R. fruticosus* showed its highest (*SM 3.2, Fig. SM.5, C*) in 2029 & 2050; indicating a reshuffling of the woody flora community.

**Fig. 5.**
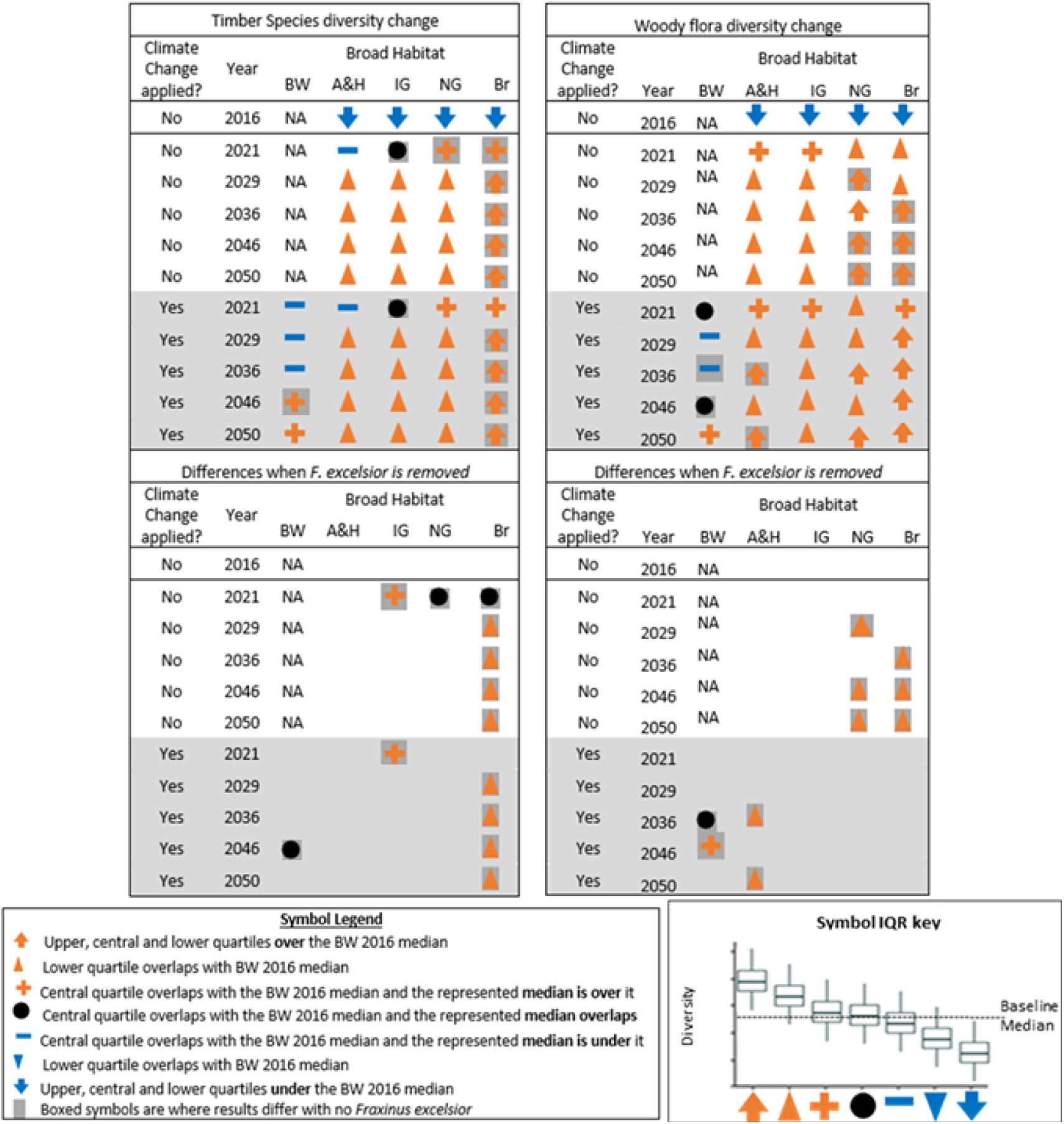
Timber and woody species group diversity change. Symbols define how the species group’s diversity score (interquartile range (IQR) and median) for a given year and broad habitat type compares with the Broadleaved Woodland baseline median (2016) given accumulation of woodland dark diversity, either with or without climate change. Thus, relative to the baseline, blue symbols indicate a decrease, black, no change and orange an increase. Broad Habitat types: BW = Broadleaved woodland (2016 baseline); A&H = Arable and horticulture; IG = Improved grassland; NG = Neutral grassland; Br = Bracken. The data used to create these boxplots was generated using an ecological niche model MultiMOVE, inputs were altered to represent baseline (1981-2016) and future climates using downscale UKCP18 climate data, incremental increase of cover-weighted canopy height representing tree growth and generalised linear mixed effect models of soil variable change under broadleaved plantation. *See, SM 3.2, Fig. SM.9 & Fig. B.10* for boxplots showing the IQRs of each species groups diversity scores per habitat and scenario.

#### 3.3.3. Impacts of removing Fraxinus excelsior

Results for the subset of data that only includes sites where *Fraxinus excelsior* habitat suitability scores suggest it is likely to be present (*see, SM 3.1*) are shown in Fig. 6 & 7. The species results (Fig. 5) build confidence in the modelling as a plausible range of common woodland trees, shrubs and herbs are predicted to have the highest modelled suitabilities. Overall there is little difference in the top species’ identity with, or without, climate change. The results for the *F. excelsior* plots (Fig. 6) estimate which species are most suitable to the newly-modelled soil and climate conditions (see more species in, *SM 3.1, Table SM.2*). The years included are 2016 (baseline,) 2036, 2046; and the net zero target year (2050).

**Fig. 6.**
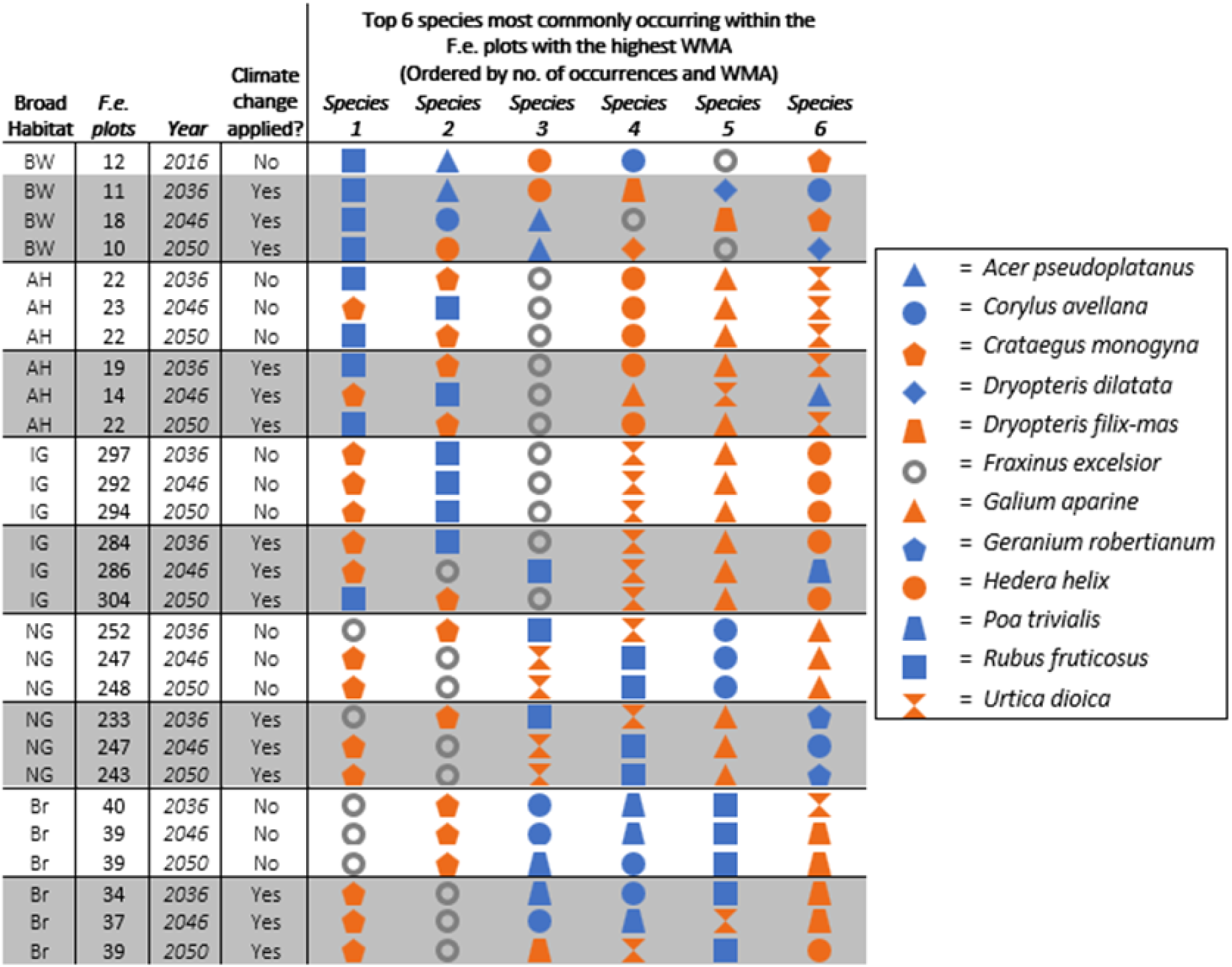
For each habitat and year with and without climate change the ***top 6 species*** most suitably predicted species are shown (top 5 if *Fraxinus excelsior* is removed). This only includes plots where weighted model average score suggests *F. excelsior* is more likely to be present than absent, determined by logistic regression. The first 2016 row represents modelled habitat suitabilities in broadleaved woodlands in the baseline year 2016 with all other years representing canopy growth and climate change scenarios. Broad Habitats: BW= Broadleaved woodland baseline; AH= Arable & Horticulture; IG= Improved Grassland; NG= Neutral Grassland; Br= Bracken; Climate change applied?: No = Baseline climate (white); Yes = predicted climate (grey). *See, SM 3.1, Table SM.2* for the top 20 species.

Within baseline broadleaved woodland, *A. pseudoplatanus* is consistently predicted as the likely replacement for *F. excelsior* under climate change as it has the highest suitability scores (Fig. 6, BW, Climate change scenarios). The vegetation results suggest little change in vegetation type with *F. excelsior* removed (Fig. 7). However, on neutral, base-rich woodlands or acid woodlands (the high-fertility habitats) the predicted forest type is low canopy woodland or scrub with species such as *Corylus avellana, Crataegus monogyna, and R. fruiticosus* (Fig. 6).

**Fig. 7.**
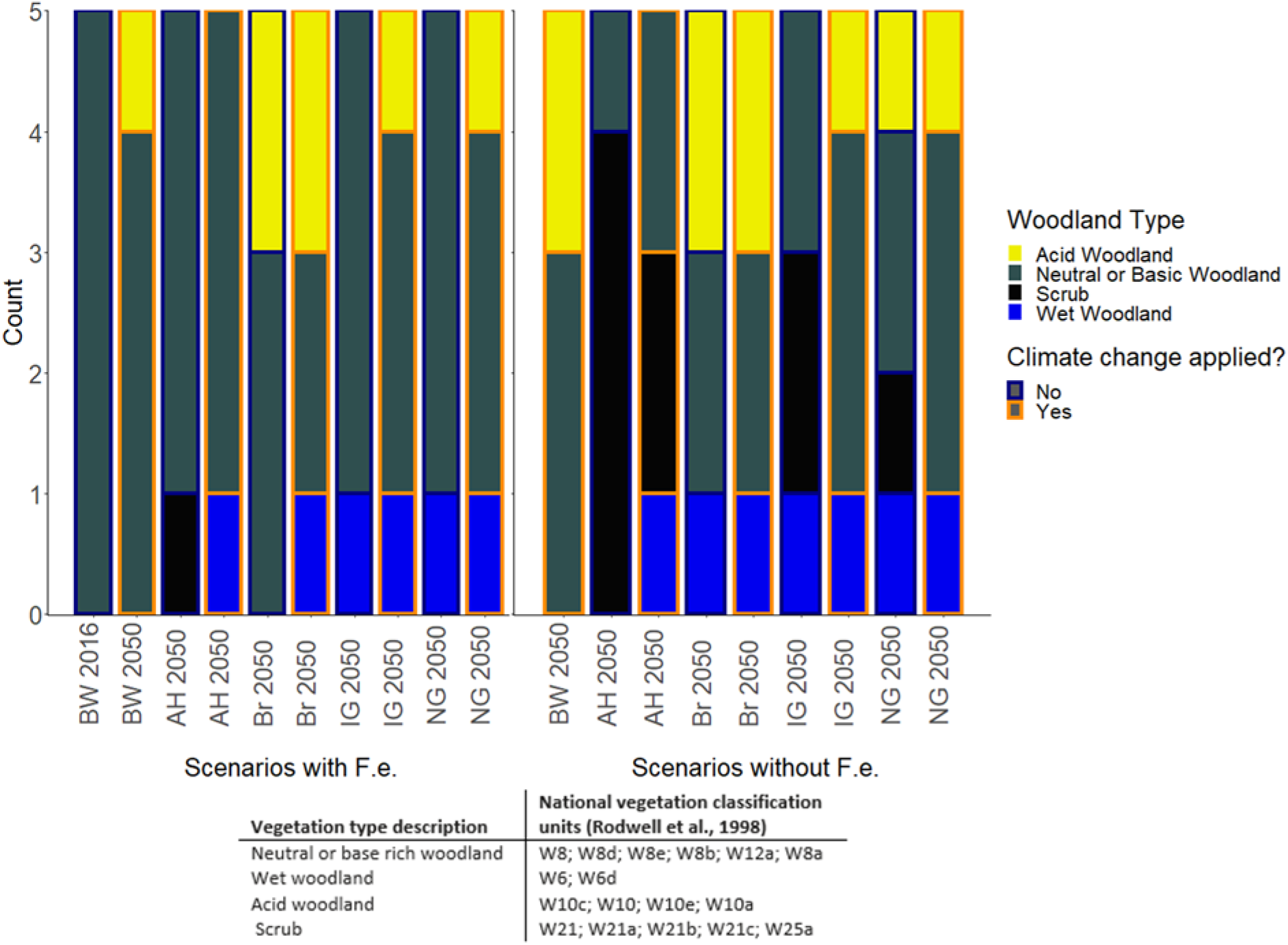
The top five woodland vegetation types for each habitat with Broadleaved Woodland 2016 baseline on the far left for reference, and 2050 for all other scenarios. The left-hand plot shows the predicted vegetation with *Fraxinus excelsior* (F.e.) within the species pool and the right plot without F.e., no baseline is shown in the right-hand plot as F.e. is present at the baseline. Woodland types were constructed via National Vegetation Classification units (NVC units, matched using the software MAVIS). *See, SM 3.1, Table SM.2,* for full data and vegetation group details. The first 2016 row represents broadleaved woodlands at baseline in 2016 with all other years representing canopy growth and climate change scenarios. Broad Habitat: BW= Broadleaved woodland baseline; AH= Arable & Horticulture; IG= Improved Grassland; NG= Neutral Grassland; Br= Bracken.

Results for improved grassland (Fig. 6, IG) also show that *C. monogyna* and *R. fruticosus* are consistently most suitable. Neutral grassland results in Fig. 6 are similar to the improved grassland but lack the higher scores of the high fertility species (*e.g. Galium aparine*). Neutral grasslands along with bracken also show the largest number of vegetation types in the top vegetation groups (*SM 3.1, Table SM.2*). An interesting feature of projected outcomes for the bracken habitat is the positive impact on fern species including *Pteridium aquilinum, Dryopteris dilatata, Dryopteris filix-mas*. These consistently occur within the top six species regardless of climate.

## 4.0. Discussion

Models can be usefully applied to manage expectations about timescales for achieving conditions suitable for understorey community assembly. This is important because understories contribute significantly to forest’s biodiversity (Brunet et al., 2012; Perring et al., 2020). Our novel inclusion of a measured local species pool allows for compositional turnover to arise as the woodland develops in the form of species changing rank based on their habitat suitability. Hence by modelling habitat suitability for a suite of potential colonists we can estimate dark diversity (potential species that could occur) at each location over time (Pärtel et al., 2011).

### 4.1 Did modelled soil and canopy change move plantation forest vegetation toward establish forest conditions?

The first aim of our modelling research was to determine if environmental conditions altered by broadleaved planting scenarios moved vegetation toward baseline broadleaved forest conditions within 30 years. Within the soil response to afforestation mixed results are shown reflecting development from differing starting conditions (Minasny et al., 2017; Mayer et al., 2020). Our results appear plausible over the timescales examined matching research into similar conditions and ecosystems (Poulton et al., 2003; Thomaes *et al.,* 2012).

The three modelled soil variables showed different trajectories. Soil pH started higher than baseline broadleaved woodland in all habitats except the less intensively-managed bracken-dominated broad habitat type (Seaton *et al.,* 2020) and then increased steadily reflecting adjustment (applied in, *2.13.2*) due to recovery from historic acid from deposition (Emmett et al., 2010).

In contrast, for nitrogen all habitats except arable were within the baseline range of broadleaved woodland values, probably because arable soils lose organic matter due to continued crop harvesting (Jones et al., 2013). However, the arable habitat was predicted to accumulate the most N, being significantly greater than baseline by 2056. But following the changes in C & N across Fig. 3, the C:N ratio is predicted to increase across the time-period. This increasing C:N over time will filter for species typical of lower fertility than in intensively-managed baseline habitats. In summary, consistent and substantial increases in C:N ratio from increase in soil C up to 2050 outweigh increases in soil N. The increase in soil C provides a positive message for forest utilisation to achieve 2050 goals in UK government strategy (HM Government, 2021).

While soil conditions are often discussed in the context of climate change mitigation (De Graff et al., 2006; Mayer et al., 2020), changing conditions also imply change in plant biodiversity and species composition. Moreover land-use legacy effects (e.g. higher N in the more intensively managed habitats) may also result in long-lasting constraints on future woodland development (Dupouey et al., 2002; Valtinat, Bruun and Brunet, 2008; Diedhiou et al., 2009).

#### 4.1.1. Importance of the starting conditions

Habitat creation can achieve large changes in abiotic conditions depending on starting conditions, but these may not be enough to achieve restoration targets. For example, switching cultivated land poor in carbon (e.g. Fig. 3, B, A&H) to perennial-dominated vegetation (e.g. Fig. 3, B, Br) can drive large changes in soil and vegetation over short time periods but still remain short of achieving target habitats (Carnus et al., 2006). In this sense magnitude of managed change from baseline will often not correlate with amount of habitat successfully created. This is representative of ecosystem function and service trade-offs (Linney et al., 2020). While plantations often have lower biodiversity than natural forests biodiversity gains from pre- to post-plantation state are still beneficial (Stephens and Wagner, 2007; Carnus et al., 2006).

Our cultivated land example (A&H) started with the lowest carbon and nitrogen values at baseline, but, made the largest gains by 2046. Although the lowest predicted increases in plant diversity of all the habitats are predicted cultivated land here (Fig. 4 & 5). However legacy effects of low carbon and high residual fertility is expected to limit the absolute biodiversity change relative to other habitat types (Mayer *et al.,* 2020; Berdeni, Williams and Dowers, 2021; Carnus *et al.,* 2006). Such legacy effects can push community assembly away from desirable endpoints (Falkengren-Grerup, ten Brink and Brunet, 2006).

In contrast, planted bracken habitats (perennial-dominated) exhibited the greatest predicted forest creation both in terms of progression toward baseline broadleaved woodland soil conditions (Fig. 3) and predicted dark diversity of forest specialist plants (*SM 3.2, Fig. SM.7, B*). Here the similarities of the broad habitats at baseline have led to forest soils and ancient woodland indicator species resembling the described target broad habitat closely. Bracken is considered as a woodland species that can become dominant after tree clearance, and there are often woodland species present as relict understories or in the seedbanks (Marrs and Watt, 2006).

Results for individual species highlight the likely importance of differences in legacy fertility in constraining community assembly (Valtinat *et al.,* 2008). The main example being improved grassland, which reflects its high residual fertility, where *Urtica dioica* is followed by *Galium aparine* (*for example in: SM 3.1, Table SM.2*). This high residual fertility on ex-agricultural land facilitates strong and rapid space pre-emption by perennial grasses and *Rubus* spp. which quickly dominate when agricultural management ceases. Therefore, future dispersal and establishment processes need to facilitate the modelled dark diversity species communities becoming realised in the future. This highlights the likely need for introduction of young trees and shrubs via planting (Ashwood et al., 2019) or a high certainty of dispersal from local populations (Brunet et al., 2012) .

### 4.2. Which trees and shrubs could replace F. excelsior with and without alteration of climate?

For the third research aim our results emphasise the importance of *F. excelsior* in UK forests and concern the potential loss of mature specimens (Mitchell et al., 2016; Skovsgaard et al., 2017; Carroll, 2020); young *F. excelsior* trees are also likely to go as soon as they grow, restricting replacement (Skovsgaard et al., 2017). The species that are predicted as the most likely replacements into the future are: *Rubus fruticosus*, *Crataegus monogyna*, *Corylus avellana*, *Hedera helix*, *Urtica dioica* and *Galium aparine*). These results are consistent Mitchell et al. (2016) suggesting that within the next 20-30 years *F. excelsior*-dominant forest is likely to remain the same community type or see an increase in acid woodland (NVC W10 scrub/low-canopy woodland communities, Rodwell, 1998) by 2050. Mitchell et al. (2016) also determined that W10 and W8 sub communities were the closest ecological analogue to British *F. excelsior*-dominated forest (*see, SM 3.1, Table SM.2*).

*Acer pseudoplatanus* is often an expected beneficiary of *H. fraxineus* (Mitchell et al., 2016; Skovsgaard et al., 2017) but is infrequent in the top six species predictions here, except where it is already established. Alternatively, if planted by land managers it is likely to establish in grasslands as it occurs in the top 10 species for improved and neutral grasslands results, making it a likely high canopy tree replacement. However, *Acer pseudoplatanus* casts a heavier shade than *F. excelsior*, therefore, the understory ground flora diversity may reduce from current levels (Mitchell et al., 2016).

The management response to ash-dieback will naturally be a determining factor in vegetation community realisation as this is effected by the disturbance regime (felling, grazing etc.) Mitchell et al., 2016 suggests. Our modelling of dark diversity suggests that planting a range of tree species suited to *F. excelsior* sites will allow for rapid re establishment of high canopy forests rather than scrub. This implies that under the starting conditions modelled here change may lead to vacant niche space; suggesting that higher biodiversity of tall canopy tree species gives ecosystems more resilience to change.

### 4.3. To what extent are these modelled assemblages changed in their species composition by projected climate change?

Climate filtering effects are noticeable for both species and vegetation. As predicted temperatures move outside of the range of MultiMOVE model’s training data (2046 onwards) after this period predictions become less reliable (*see*, *SM 4, Fig. SM.10*). However, the range of the climate training data for MultiMOVE is large given that it covered the full extent of Great Britain (South-East-England to Scottish mountains). Modelling the ecological impacts is an acknowledged challenge and even dynamic approaches need to estimate the uncertain consequences of no-analogue climate space and novel competitive interactions (Williams and Jackson, 2007; Mouquet et al., 2015; Alexander et al., 2016). *See, SM 4.2,* for further details on modelling into future climate space.

Nevertheless, our results showed that none of the scenarios of future afforestation are likely to replicate vegetation assemblages typical of baseline woodlands in our Welsh study area. This implies that expectations require careful management and that targets based on the present or past may need to be applied loosely to future forest development.

The highest diversity is within the Timber and Woody species groups, which is expected as the model workflow is deliberately set up to represent forest establishment, building confidence in its robustness. Comparisons within our results, however, highlight novel differences under different climates, habitats, and where *F. excelsior* is removed assuming *H. fraxineus* induced-mortality. The baseline climate scenario shows a steady increase in Timber species diversity across habitats (*SM 3.2, Fig. SM.9*); with the residual fertility of the grassland starting habitats being the likely cause of their higher scores; this is exacerbated by increasing soil pH which is recovering from sulphur deposition. The lowest scores for timber species coincide with higher temperatures in the predicted climate data (*SM 4, Fig. SM.10*); this is assumed to be linked to drought stress, already recognised as a concern for UK native timber species (Broadmeadow *et al.,* 2005). As *F. excelsior* along with *Fagus sylvatica* and *Acer pseudoplatanus* are predicted to perform better with climate change scenarios (Broadmeadow *et al.,* 2005) its potential loss is a major concern.

For the Woody flora group, with and without *F. excelsior,* the planting scenario scores differ much less than under baseline and predicted climate change scenarios for Timber species (*SM 3.2: Fig. SM.10*, *B&D* varies more than *Fig. SM.9 B&D*). Thus, climate may have less of an effect on the dark diversity of the totality of trees and shrubs in the species pool (e.g. Broadmeadow et al., 2005; Hastings et al., 2014). Recent empirical work across Europe matches this, showing increases in shrubby flora due to climate change (Perring et al., 2020).

Modelled Nectar plants and AWI species diversity (Fig. 4) remained below the baseline broadleaved woodland reference in all habitats throughout the modelled period modelled. The implication is that despite gradual change in soil conditions resulting from forest development, legacy differences in soils persist and inhibit change toward conditions more typical of baseline forest (Valtinat *et al.,* 2008). The few instances where predicted habitat suitability exceeds the baseline is in the later years with climate change applied. This is especially notable in the Nectar plants where broadleaved woodland modelled under a predicted climate shows higher scores than baseline in 2050. This result suggests that Nectar plants are favoured by predicted climates in 2050. This is notable in results for individual species for example *Rubus fruticosus* (*SM 3.2, Fig. SM.5, C*); *Hyacinthoides non-scripta* (*SM 3.2, Fig. SM.4, C*) and *Crataegus monogyna* (*SM 3.2, Fig. SM.5, A*).

## 5.0 Conclusions and implications

The plant species and vegetation modelling suggest that woodland with *F. excelsior* absent is most likely to establish as scrub or low canopy woodland within the next 30 years. Alternatively, where it is present in the local area within established woodlands, *A. pseudoplatanus* will become the main replacement. While *F. excelsior* dominated vegetation is likely to shift substantially, new species compositions do not appear likely to emerge, even under predicted climate up to 2050. Likely new replacement dominants (other than *A. pseudoplatanus*) being *Crataegus monogyna*, *Corylus avellana*, *Rubus fruticosus* and in some circumstances *Pteridium aquilinum* and *Dryopteris* spp.; any changes are likely to take a decadal timeframe to be observed in woodland communities given successional timescales. This may also be prolonged as the removal of *F. excelsior* from the species pool used here is a simplification of reality more likely to be a gradual decline (*see, SM5.1, for details*). Given the fact that the UK already has extremely fragmented and degraded habitats (Watson et al., 2011; Hayhow et al., 2019; Forest Research, 2020) it seems unwise to abandon them to already degraded successional processes under the assumption they will sort themselves out.

As plantation scenarios do not show consistent convergence with the baseline across all the species and soil variables; we cannot say definitively that plantation can establish to equivalent baseline broadleaved woodland conditions even after 30 years. However, as the baseline woodlands are likely to be hundreds-of-years-old (The Woodland Trust, 2020) complete congruence would be unlikely, particularly for Ancient Woodland Indicators. Our results do show some convergence with broadleaved woodland within 20 years for soils and some habitats (Bracken, Improved & Neutral grassland’s); showing overlapping herbaceous species group diversity in their upper ranges from the late 2020s or 2030s onwards. Determining sites to achieve forest establishment within less than successional timeframes (30-40 years) would be highly beneficial for carbon goals thus is an important avenue for further research, as availability of land for planting as well as suitability of land is important for forest creation.

Suggestions for management and policy derived from these results and synthesis from review of other publications can be found in *SM5.2*.

## Supporting information

Table SM.1. afforestation_soils_change

Table SM.2. Top_20Sp._in_Fraxinus_excelsior_plots_and_NVCunits

Table SM.3. MultiMOVE and forest ecosystem functions or services species list

Neural_net_building_files_V2.0

## Acknowledgements

This work was funded by:

- NERC Envision Doctoral Training Programme
- Environment & Rural Affairs Monitoring and Modelling Programme (ERAMMP) (Welsh Government Contract C210/2016/2017)
- UK-SCaPE program delivering National Capability (NE/R016429/1)

We thank the Welsh Government and the ERAMMP team for data provision and the downscaled projected future high emissions (RCP8.5) climate data were derived from UKCP18 Regional Climate Model projections produced as part of the UK-SCAPE programme (Natural Environment Research Council, NE/R016429/1). Lucy Skelton and Nicola Spence (Department for Environment, Food and Rural Affairs); Jo Clark (Future Trees Trust); Clare Trivedi and the UK conservation team at the Millennium Seed Bank (Royal Botanic Gardens Kew) provided guidance on literature and Timothy Makower (Lancaster University) assisted with the literature review.

Lastly, BW thanks the late Jonathan West for inspiration and guidance that led me to pursue this work.

## Authors’ contributions

BW and SS conceived the initial ideas, BW constructed most of the modelling workflow with the calibration neural networks created by SS. AK, RM, SK and ER contributed data or helped with data acquisition. BW created the original manuscript with subsequent editing and comment from all authors.

## Data availability statement

Summarised modelling data can be found within the supplementary material and baseline datasets are within: Smart et al., (2020); Maskell at al., (2020); Robinson et al., (2022).

## Supporting Material (SM) and data

### SM 1 | Description of Broad Habitat Types

The six broad habitats are described in Jackson (2000). During the baseline Glastir Monitoring and Evaluation Programme (GMEP) survey areas of land were assigned to these six broad habitats among others using a vegetation key available online at: http://nora.nerc.ac.uk/id/eprint/5194/1/N005194CR.pdf.

The top 10 most common species in quadrats surveyed in GMEP and assigned to each broad habitat were as follows:

- **Broadleaved woodland (BW)**, Broadleaved, Mixed and Yew Woodlands being the full name: Rubus fruticosus, Dryopteris dilatata, Hedera helix, Fraxinus excelsior, Thuidium tamariscinum, Corylus avellana, Hyacinthoides non-scripta, Dryopteris filix-mas, Sorbus aucuparia, Pteridium aquilinum.
- **Coniferous Woodland (CW):** Picea sitchensis, Dryopteris dilatata, Vaccinium myrtillus, Thuidium tamariscinum, Plagiothecium undulatum, Mnium hornum, Rubus fruticosus, Rhytidiadelphus loreus, Deschampsia flexuosa, Sorbus aucuparia.
- **Arable and Horticulture (A&H):** Poa annua, Poa trivialis, Ranunculus repens, Trifolium repens, Lolium perenne, Agrostis stolonifera, Taraxacum agg., Persicaria maculosa, Rumex obtusifolius, Triticum aestivum.
- **Improved grassland (IG):** Lolium perenne, Trifolium repens, Ranunculus repens, Holcus lanatus, Cerastium fontanum, Poa trivialis, Taraxacum agg., Agrostis capillaris, Poa annua, Rumex obtusifolius.
- **Neutral grassland (NG):** Holcus lanatus, Agrostis capillaris, Trifolium repens, Lolium perenne, Ranunculus repens, Cerastium fontanum, Anthoxanthum odoratum, Taraxacum agg., Cynosurus cristatus, Rumex acetosa.
- **Bracken (Br):** Pteridium aquilinum, Agrostis capillaris, Rhytidiadelphus squarrosus, Pseudoscleropodium purum, Anthoxanthum odoratum, Galium saxatile, Holcus lanatus, Festuca ovina, Potentilla erecta, Pleurozium schreberi.

## SM 2 | Modelling change in soil variables under afforestation

Determining how soils change in response to afforestation via planting or natural succession was completed by constructing generalized linear mixed-effects models trained on data gathered from a review of the literature where the effects of afforestation on soil variables had been measured under time series or chronosequences. The use of biogeochemistry models to dynamically process the impact of afforestation was not adopted because the necessary soil measurements do not exist for each modelled location. Using average inputs coarsely resolved to larger grid squares and dominant soil type would have greatly decreased the accuracy and realism of the modelling process removing the benefit of filtering the observed plant species composition at our very high-resolution sampling sites.

The four soil variables (C, N, pH, moisture) were modelled independently of each other as an insufficient number of studies measured all the variables together so there was too little data to account for relationships. Here, we assume that the modelled change is reasonably auto-correlated with other soil variables whereas in reality we know they respond as an intercorrelated complex. Thus, the modelled results are strongly dependent on the studies selected from the literature and the realism of the modelled change in each variable.

The best performing model for soil moisture change (lowest AIC, highest R^2^) consisted of forest type and starting Ellenberg wetness value, this lead to a uniform response across modelled years.

Despite data availability falling short of the representativeness and robustness we would have liked for the study habitat types, a clear benefit is that the models created here summarise changes in soil conditions that have been observed resulting from afforestation on soil and vegetation starting points that equate with the soils and vegetation of the GMEP baseline. Readers can therefore inspect the underlying data (see below) and assess the representativeness of the observations and the robustness of the estimated soil changes.

The collation of data from the literature below can be found in the file:

“Table SM.1. afforestation_soils_change.csv”.

To some extent, the lack of such long-term fundamental datasets is surprising. Despite a number of long-term experiments existing across the UK and ecological science being an endeavour that is at least 200 years old, there is a lack of fundamental information on long-term soil and vegetation changes in response to human management.

### SM 2.1 | Literature sources

Care was taken to closely match the methods of each study found during the literature review to the GMEP methods or to ensure that data values could be converted to match. Non-UK studies were omitted. This search resulted in datasets of varying size. Requests were made to study authors to provide full datasets, while we also included relevant open access data; A summary of the studies is as follows:

- A 100 year chronosequence of 40 plots in Kielder forest (Vanguelova *et al*., 2019)
- Forest conditions development under different tree species in a chronosequence (Ovington, 1953)
- A pooled selection of former agricultural sites planted up with broadleaves (Ashwood *et al*., 2019)
- Rothamsted regenerating broadleaved woodland post-abandonment from arable (Jenkinson, 1971; Poulton *et al*., 2003)
- A study of succession on lowland heaths (Mitchell *et al*., 1997)
- Plots extracted from the UK Centre for Ecology and Hydrology’s Countryside Survey data showing vegetation trends (increase in scrub, bracken or tree species cover) as having undergone afforestation (Reynolds *et al*., 2013)
- Dataset of afforestation provided by Aidan Keith from work on short rotation forestry soil development (Keith *et al*., 2015; R. L. Rowe *et al*., 2016)

### SM 2.2 | Data categorisation and model construction

Data was categorised in two ways (for: total C%, total N%, pH) in order to ensure robust fitting to scenarios, by afforestation type (planted broadleaved, planted conifer, or succession), and by taxon (gymnosperm trees, angiosperm trees, or bracken). The two different data categorisation types were used as insufficient data on nitrogen was found for it to be modelled according to its afforestation type. This categorisation was done per-treatment within each study. Where studies used chronosequences, controls e.g. grassland or arable sites, were used as time 0 matches for each variable. Also where multiple controls or multiple treatments were replicated the variables of these were averaged to become start or end values as appropriated. This was also done where samples had been taken from multiple soil depths e.g. samples taken from 0-5, 5-10 and 10-15 cm where averaged to give a 0-15 cm depth value. All soil data recovered during the literature review was taken from within the O horizon to match the baseline GMEP data which was taken from the field layer (0-15 cm).

Calculating the change in variables was done by subtracting the value at start time (or matched time 0 for chronosequence data) with the values at treatment end time and dividing by years duration of the study to give a change of X in a variable per year. This approach takes advantage of the correlation between variables within sites, between site differences were accounted for by a study site random effect. However, given the variability in the data different model constructions were used these being:

Ø pH change per year as determined by afforestation type & time are predictors
Ø Change in total carbon per year was transformed then afforestation type & value at the start were used as interacting terms alongside time as predictors
Ø Total nitrogen was the same as carbon but time & value at start were used as interacting terms alongside afforestation type

Exact model constructions can be found in the Boxes 1-4 below.

As the poorest data was found for soil field moisture% with only a few treatments with this data gathered this data was categorised by habitat type (conifer forest or broadleaved forest). Also as this data was so noisy, no good model could be made for the raw field moisture data thus the neural networks used in the model workflow to calculated Ellenberg wetness (EbF) were used to gain EbF values and these were modelled. The model construction can be found in Box 4. The planted results from the Ellenberg wetness model show no convergence (*SM.2.2,* Fig. SM.1) towards baseline woodland most likely due to planted forest holding onto underlying soil properties from the baseline habitats, thus are not explored in the main body of work.

In grassland and bracken habitats N is modelled to reach 0 or negative values for a handful of plots from 2046 onwards. These plots’ values represent where the model fails to reproduce a possible reality and are thus excluded from all the plant results.

**Figure SM.1.**
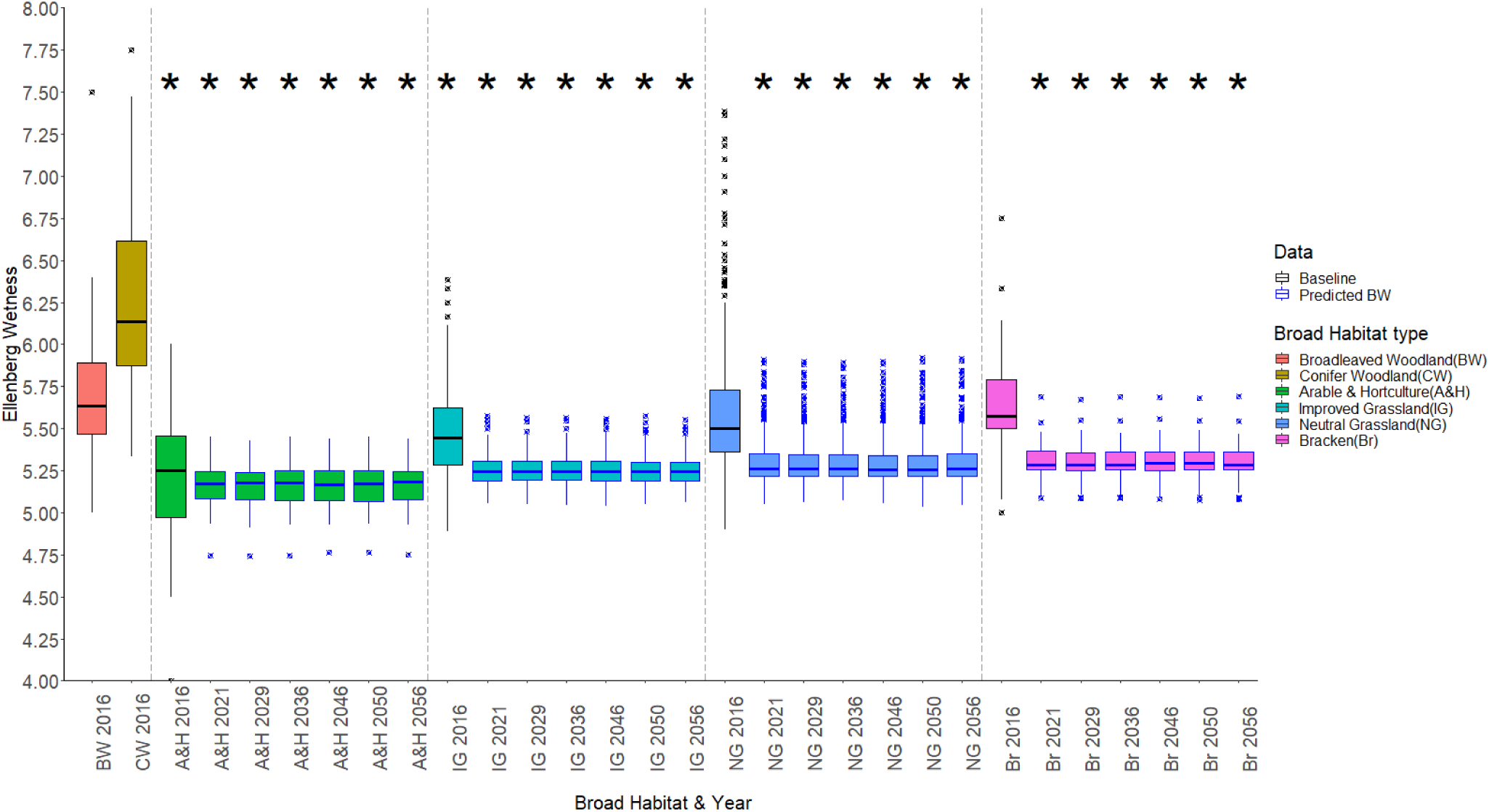
Box plots of Ellenberg Wetness per Broad Habitat (BH) type and year. The year 2016 for each BH type represents the observed data (black boxes), subsequent years data were predicted by using 2016 data as inputs to generalised linear mixed effect models of broadleaved woodland plantation (blue boxes). Asterisks (*) at Y=7.5 represent significant differences (p-value=0.05) of logged variables of the 2016 Broadleaved Woodland baseline compared to the Broad Habitat type and year below each asterisk.

### SM 2.3 | Model construction and formula code

#### Box 1.

pH model construction and summary from R. Formula: Change in pH per year ∼ afforestation type + years from start year + random effect for treatment site; planted_vs_succession = afforestation type.

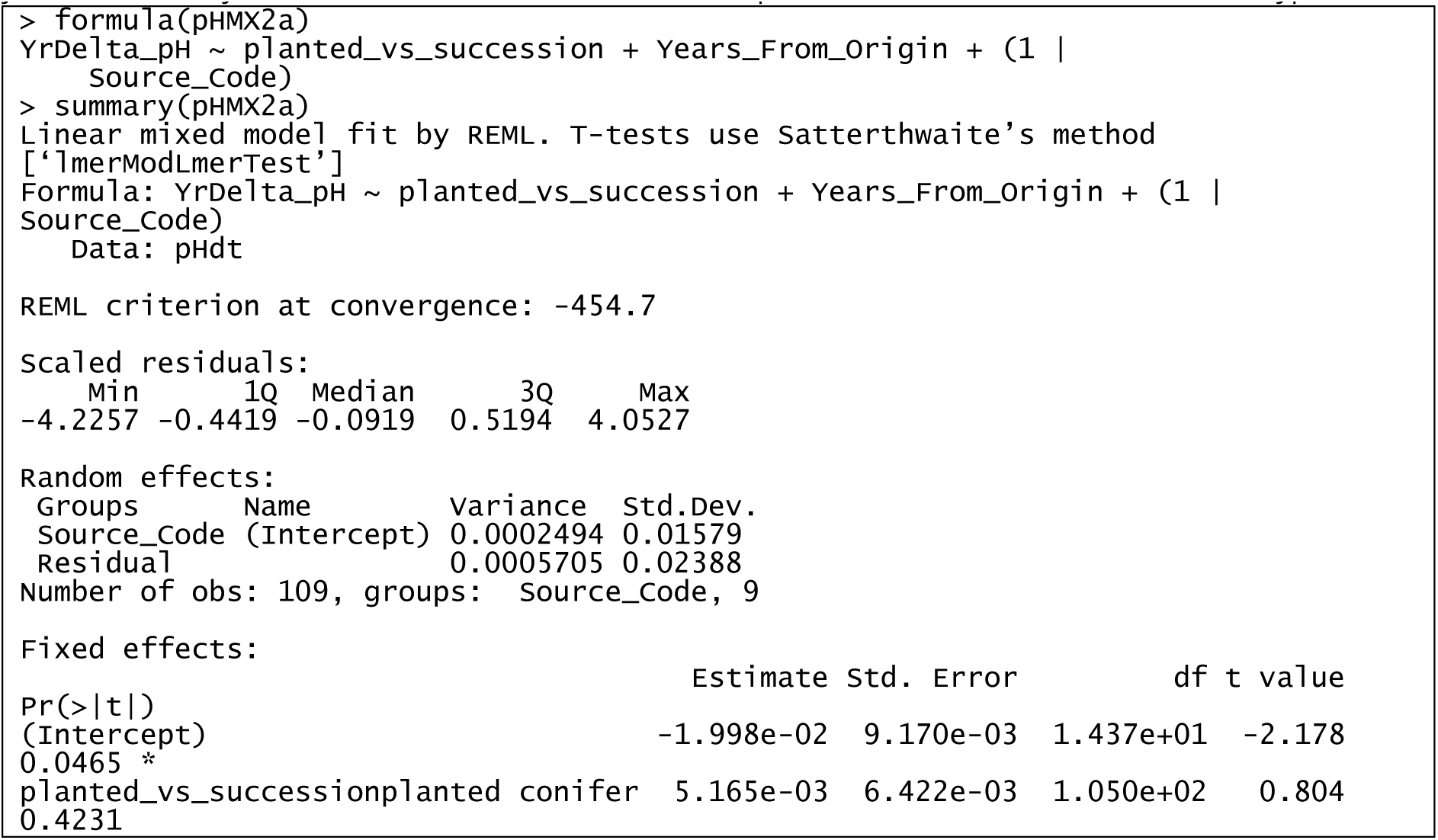
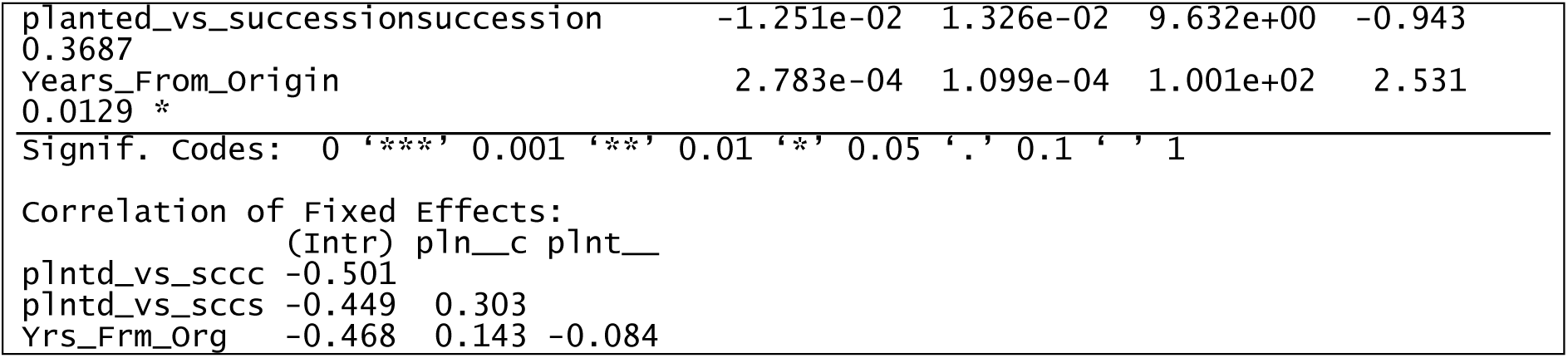

#### Box 2.

Carbon model construction and summary from R. Formula: transformed total soil carbon change per year ∼ (afforestation type + total soil carbon at start year) + years from start year + random effect for treatment site. Transformation: (X+5)^2.6^, where X is total soil carbon change per year; planted_vs_succession = afforestation type.

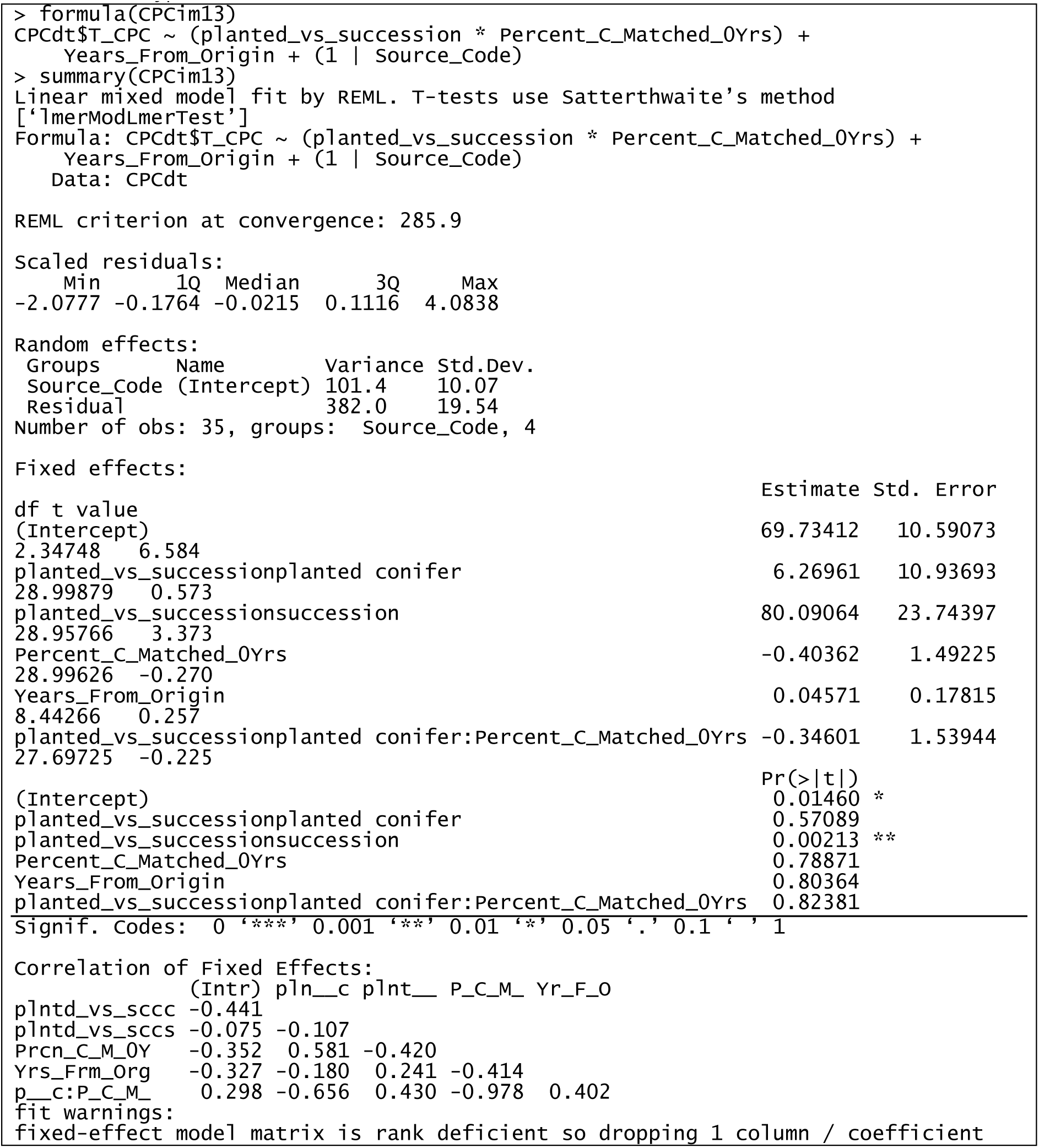

#### Box 3.

Nitrogen model construction and summary from R. Formula: transformed total soil nitrogen change per year ∼ (Years from start year * Total soil carbon at start year) + taxon + random effect for treatment site. Transformation: (X+1)^4.9^, where X is Total Soil Nitrogen change per year.

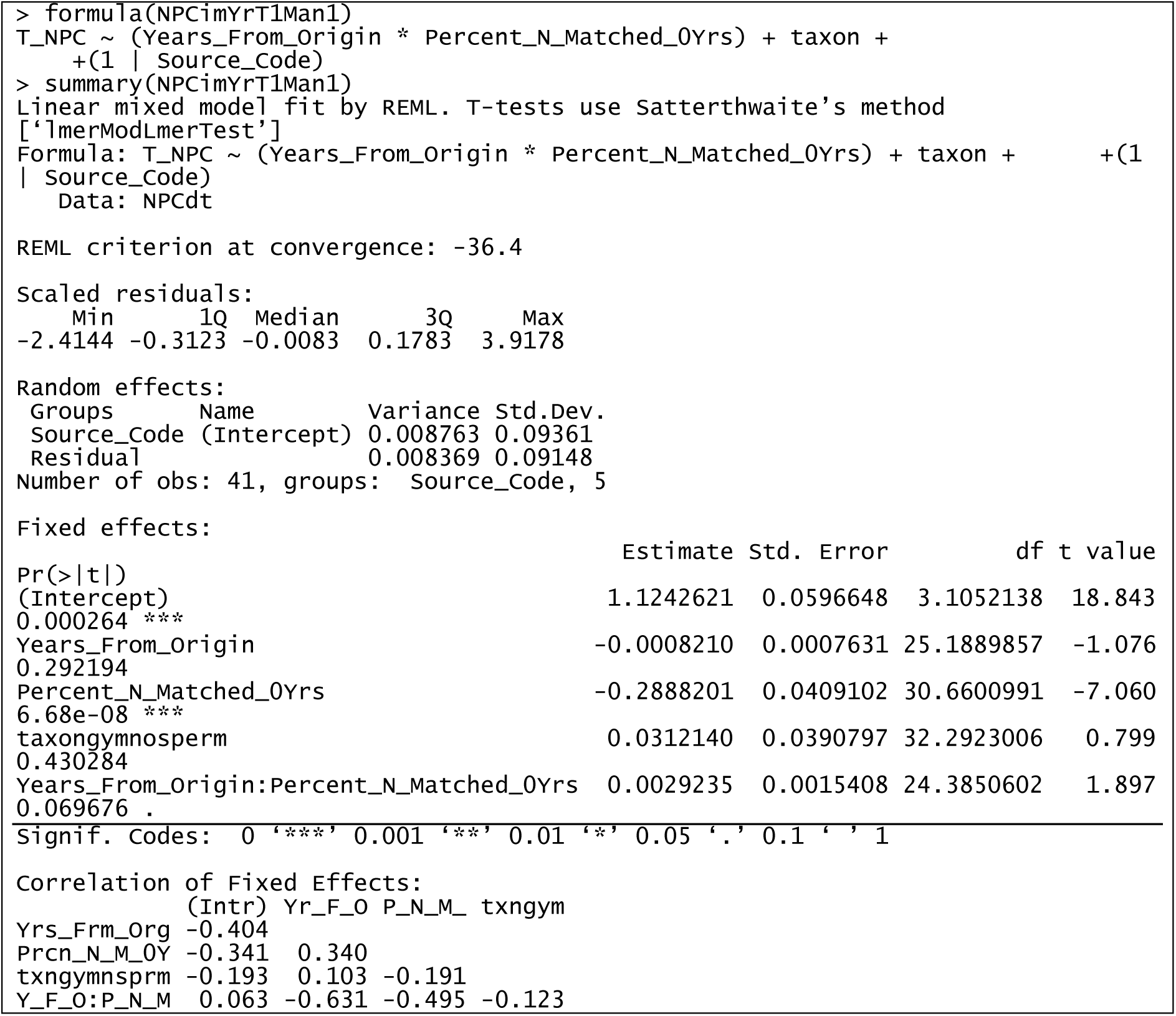

#### Box 4.

Ellenberg wetness (EbF) model construction and summary from R. Formula: over all change in EbF ∼ (Habitat type * EbF at start year) + random effect for treatment site; conifer_broadleaved_openscurb = Habitat type.

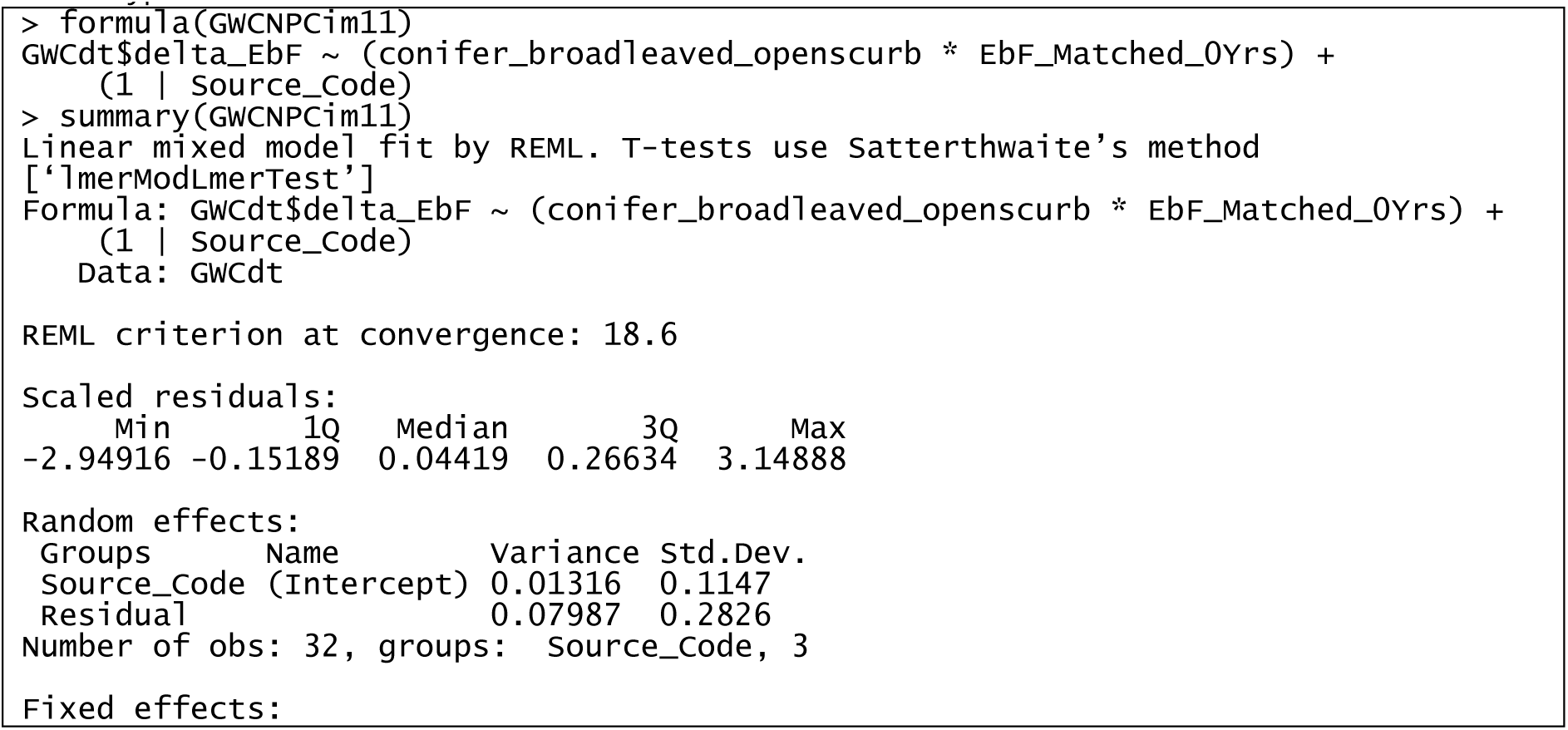
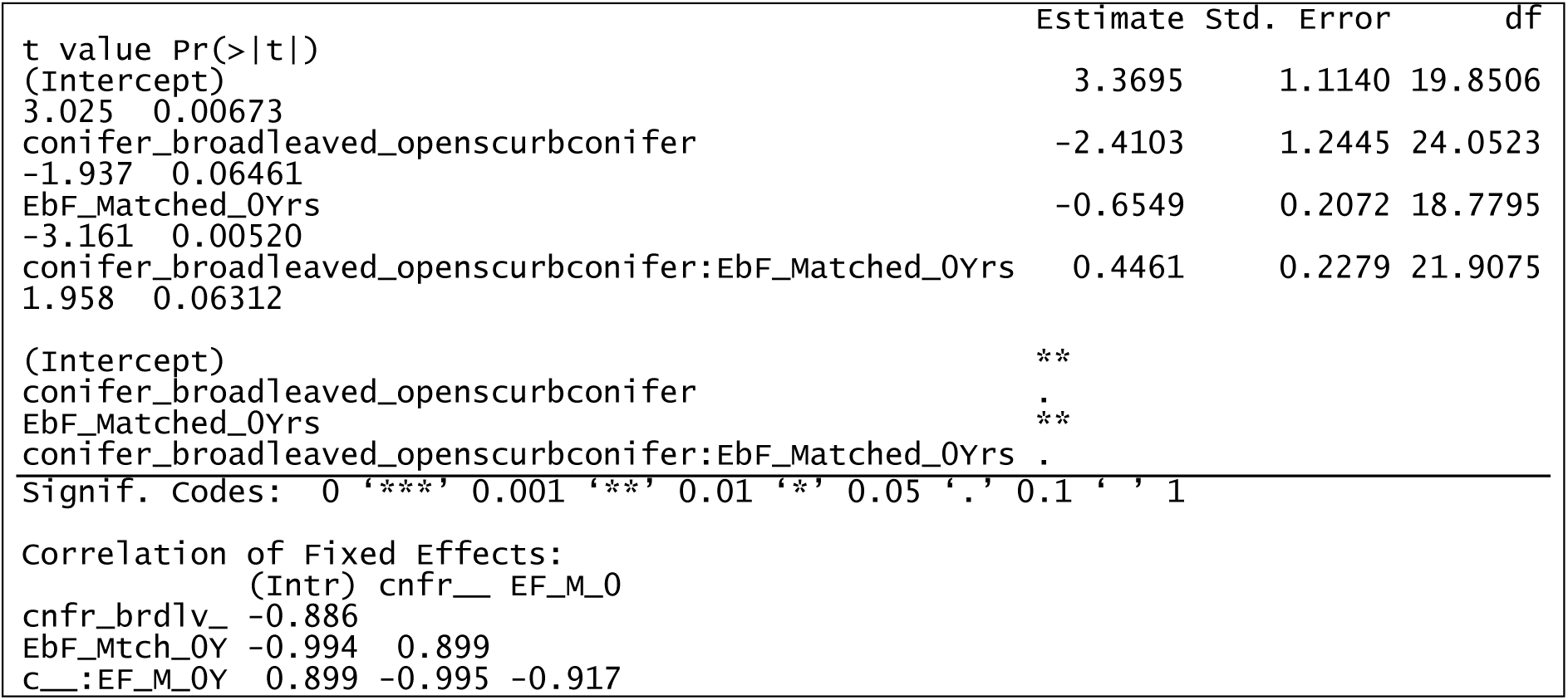

### SM 2.4 | Ellenberg indices neural net calibration

MultiMOVE accepts Ellenberg indices as inputs. These convey the ecological position of the soil and vegetation along gradients of soil moisture, fertility and pH (MultiMOVE species niche viewer Shiny app here https://shiny-apps.ceh.ac.uk/find_your_niche/). Because we were interested in how changes in soil conditions change habitat suitability for the plant assemblage, we constructed neural network models that could predict mean Ellenberg indices from measured or modelled soil variables. The latter step was achieved by constructing a model per abiotic gradient where the predictors were the soil measurements from the 5% of the training data for MultiMOVE that comprised fine resolution joint recording of soil and plant species composition. These co-located samples came from a high quality unbiased and representative sample of British vegetation types (Carey et al., 2008; Smart et al., 2010). Models were tuned, trained and tested using the neural net R package (Ripley, 1994). Predictors for each Ellenberg score gradient were standardised by their range to vary between 0 and 1 and the data split at random into 70% training and 30% testing.

The neural net method was chosen because we were interested in constructing the predictive model with the greatest accuracy but based on a small set of predictors all with strong ecological justification for their inclusion. Given the possibility of overfitting we tested the transferability of each model by comparing the predictions from the final neural nets against a similarly designed but totally separate survey of soil and vegetation carried out across Wales between 2013 and 2016.

See the zip file: “Neural_net_building_files_V2.0.zip”, for the detailed method and code and construction data.

## SM 3 | Workflow outputs and graphical plots

**Figure SM.2.**
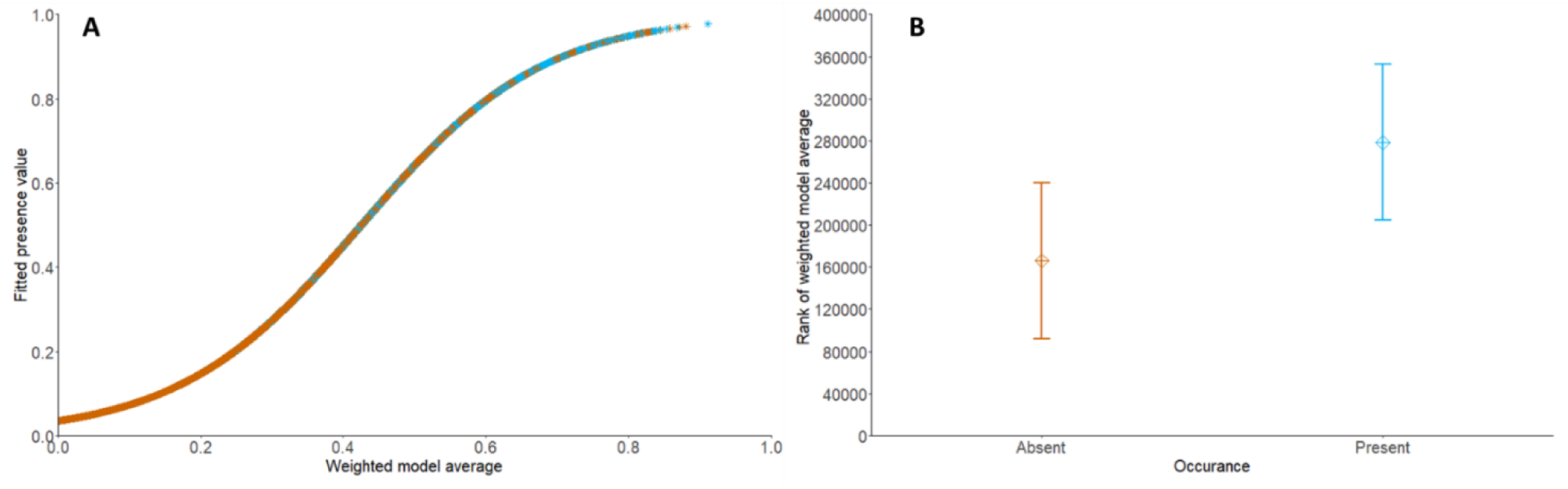
Results of testing the species ecological niche model MultiMOVE outputs against the observed baseline. A. Logistic regression (LR) of weighted model averages (WMA) of species per X-plot observed as present (light blue) or not present (brown). A species within a plot having a WMA of 0.43 or over according to the LR is more likely to be observed as present (fitted presence of 0.51) than absent within the input data (logistical regression WMA model coefficients P-value <0.001). B. Average rank plots of WMA values with rank standard deviations for absent species (brown, Absent) versus present in each modelled quadrat (light blue, Present).

Logistical regression showed that an increase in a species suitability score increases the probability of the species being observed as present in a plot within the input data (0.43 or over gives a fitted presence value of 0.51, logistical regression WMA model coefficients P-value <0.001; Fig. SM.2).

### SM 3.1 | Fraxinus excelsior plot Species and NVC units

Results within the *Fraxinus excelsior* plots and NVC units table are a subset of the total data, only including sites where *Fraxinus excelsior* habitat suitability scores (weighted model average) of 0.383 or over suggest the species is likely to be present (>0.51) according to logistic regression conducted on baseline scores.

Within *Fraxinus excelsior* plots the top 20 species by the number of plots they occur in and National Vegetation Classification (NVC, matched using the software MAVIS) units determined with and without *F. excelsior* can be found in the file:

“Table SM.2. Top_20Sp._in_*Fraxinus*_*excelsior*_plots_and_NVCunits.xlsx”.

### SM 3.2 | Species within MultiMOVE and the ecosystem function and service groups

The full species list MultiMOVE uses with the categorisation columns of the ecosystem functions or services species groups can be found within the file:

“Table SM.3. MultiMOVE and forest ecosystem functions or services species list.csv”.

**Figure SM.3.**
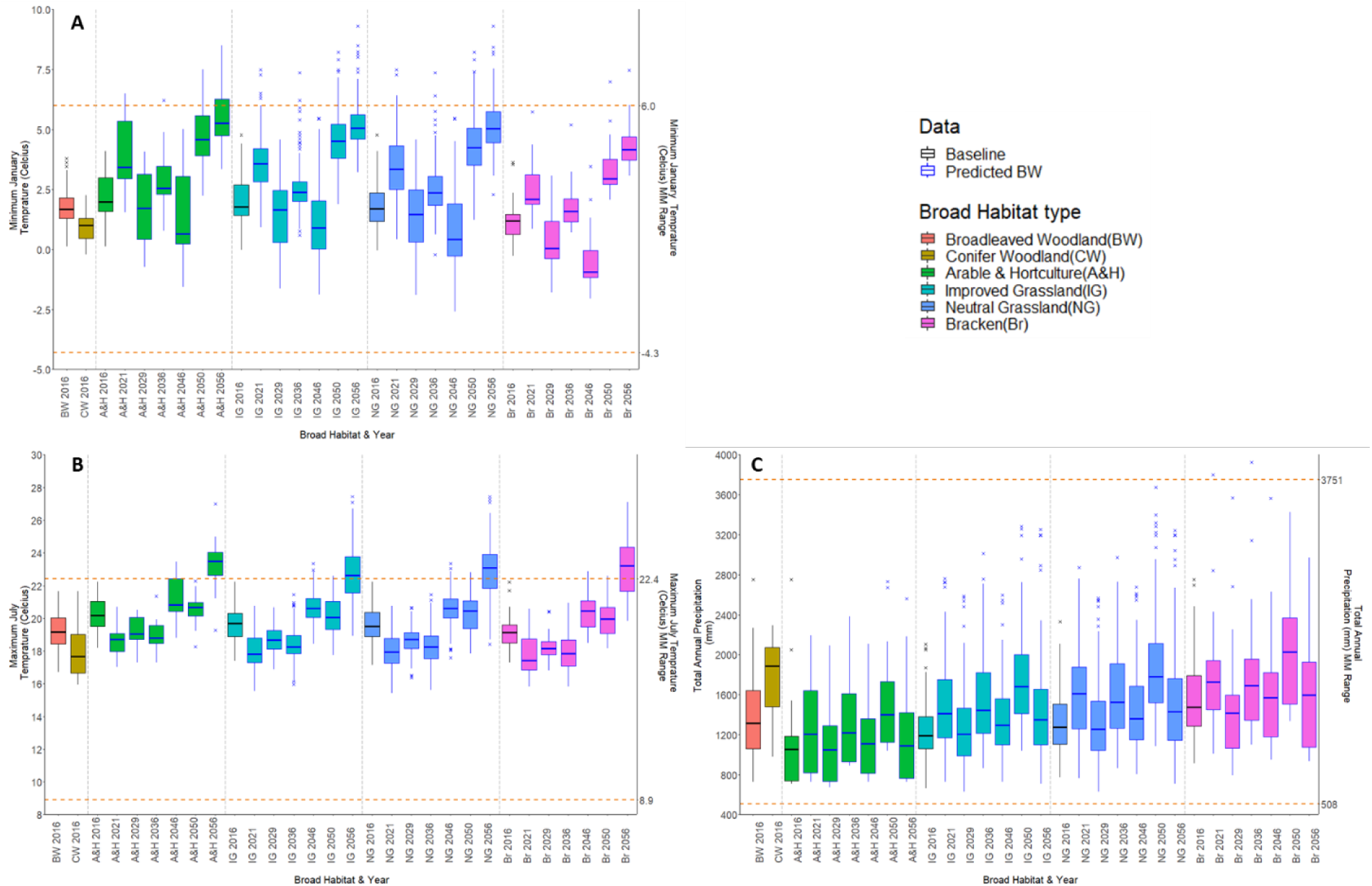
Boxplots showing Glastir Environmental Monitoring Program X-plot climate data from UKCP18 at 1 km square scale from 2016 (averaged from 1981 to 2016 as a baseline) through subsequent predicted years up to 2056. Observed (2016, black edged box plots) data was sourced from Met Office HadUK-Grid, 1 km climate data and averaged. Predictions data (blue edged box plots) from the UKCP18 high emissions scenario, RCP8.5, UK regional 12 km scale probabilistic data, was downscaled to 1km. A. Total annual precipitation in mm, B. minimum January temperature in °C and C. Maximum July temperature in °C; georeferenced to the 1 km square each X-plot was within. Left hand Y-axes shows variable ranges to derive boxplot values from (black observed average for 2016; blue beyond 2016 predicted). The dashed orange lines show the top and bottom of ranges the R package MultiMOVE was constructed within. X-axis labels show the X-plot groups of year and board habitat type: A&H = arable and horticulture; I.G. = improved grassland; N.G. = neutral grassland; Br = Bracken.

**Figure SM.4.**
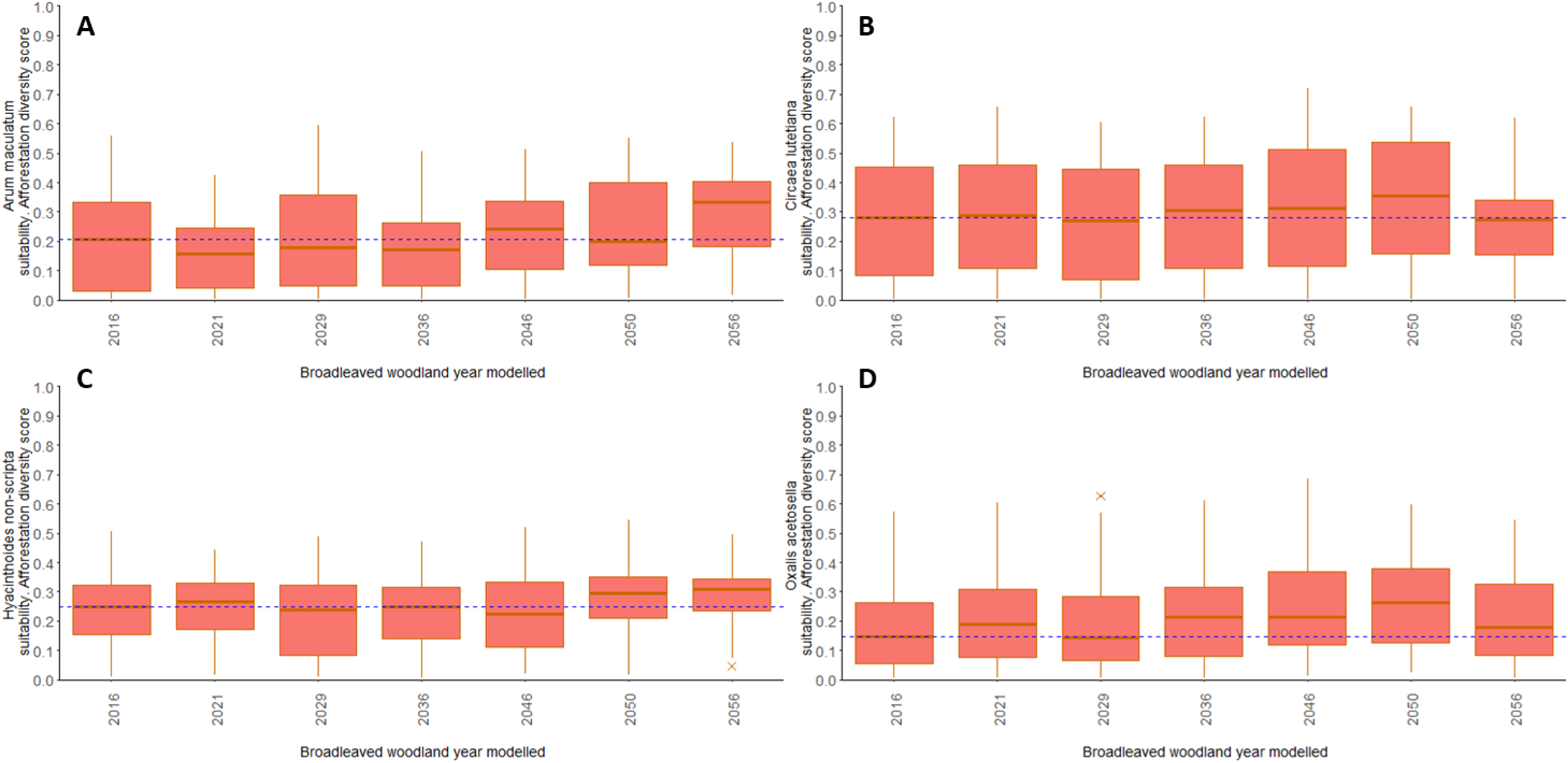
Boxplots of *Arum maculatum*(A), *Circaea lutetiana*(B), *Hyacinthoides non-scripta*(C), and *Oxalis acetosella*(D), MutliMOVE output results. Data represents broadleaved woodland at baseline in 2016 from Glastir Monitoring and Evaluation Program data under predicted climate data (UKCP18).

**Figure SM.5.**
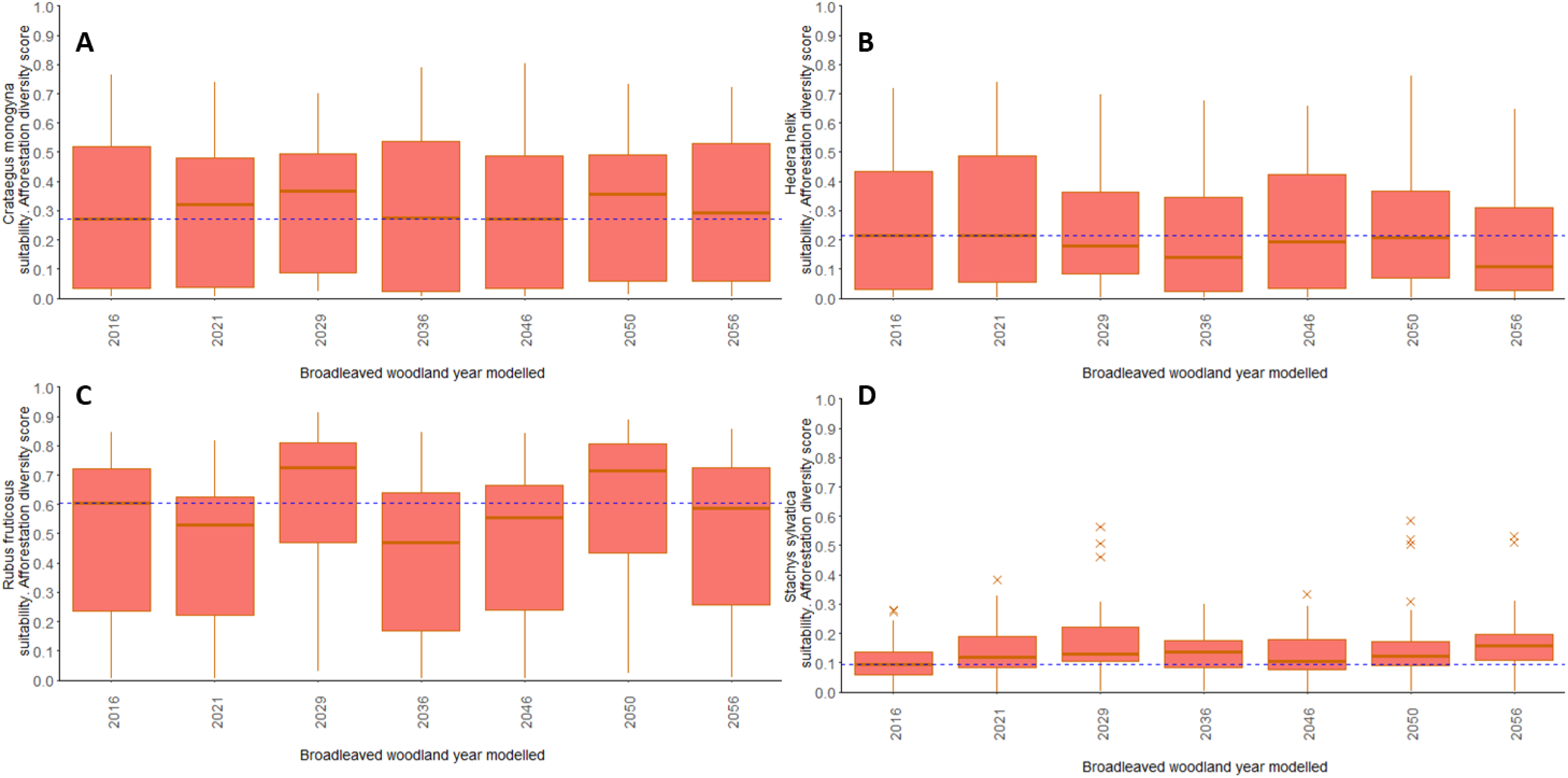
Boxplots of *Crataegus monogyna*(A), *Hedera helix*(B), *Rubus fruticosus*(C), and *Stachys sylvatica*(D), MutliMOVE output results. Data represents broadleaved woodland at baseline in 2016 from Glastir Monitoring and Evaluation Program data under predicted climate data (UKCP18).

**Figure SM.6.**
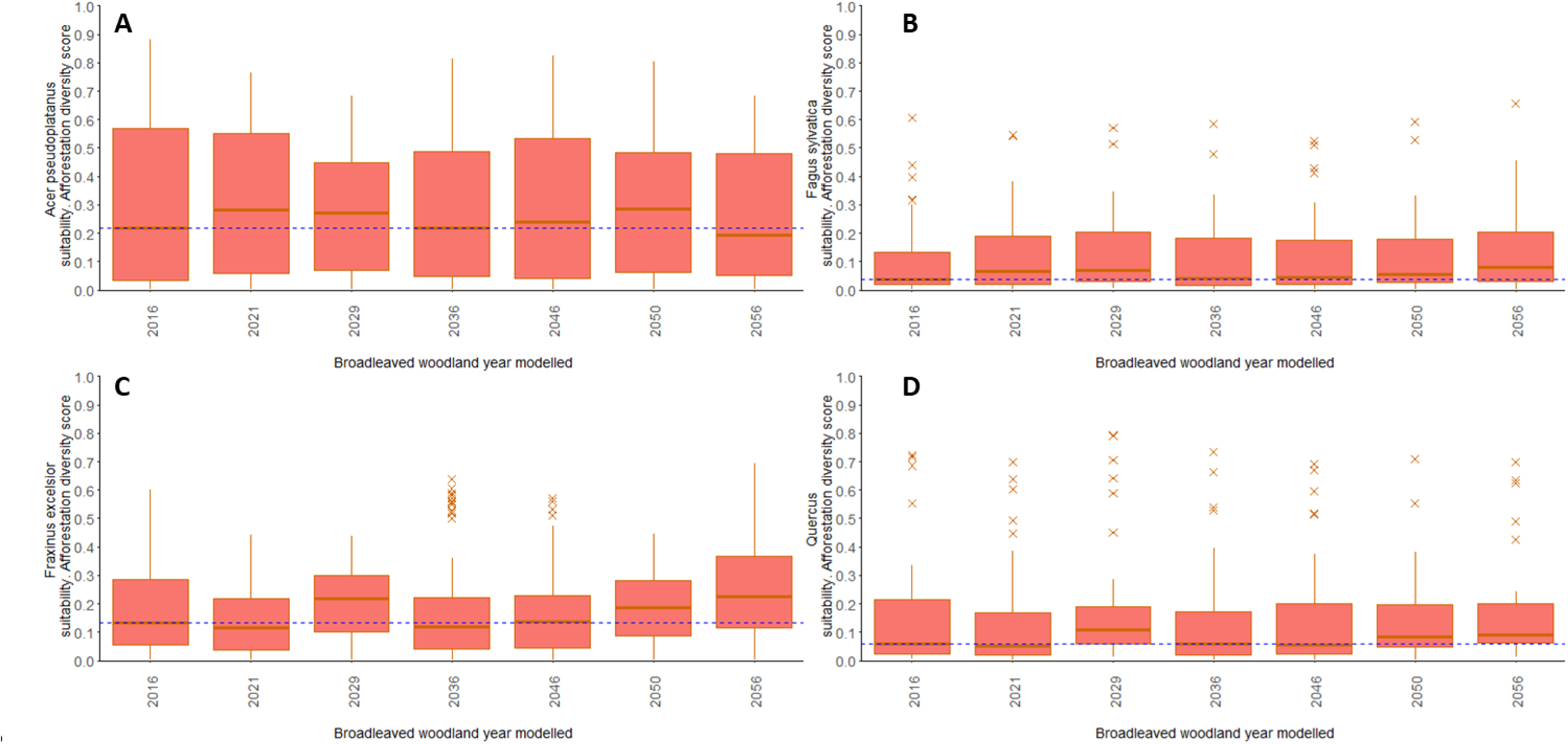
Boxplots of *Acer pseudoplatanus*(A), *Fagus sylvatica*(B), *Fraxinus excelsior*(C) and *Quercus Sp.* (D), MutliMOVE output results. Data represents broadleaved woodland at baseline in 2016 from Glastir Monitoring and Evaluation Program data under predicted climate data (UKCP18).

**Figure SM.7.**
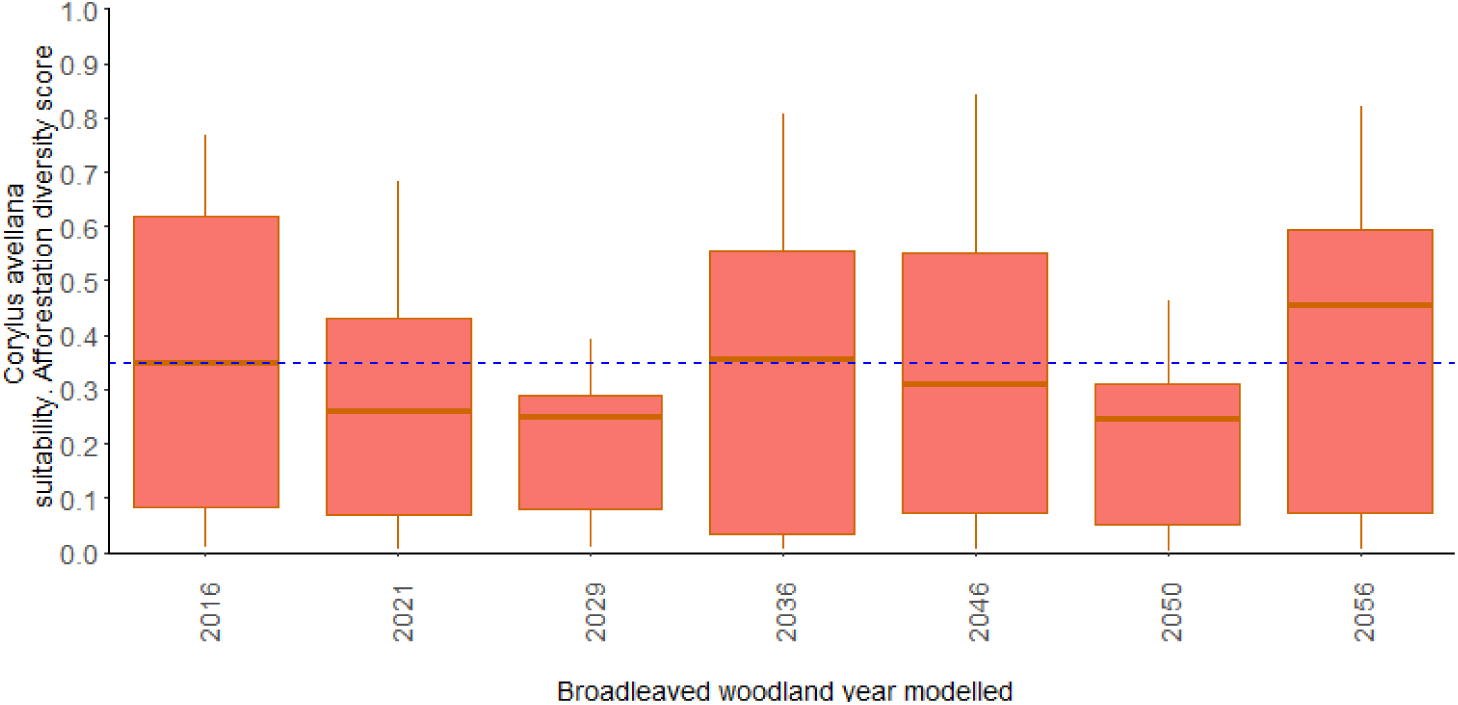
Boxplots of *Corylus avellana*, MutliMOVE output results. Data represents broadleaved woodland at baseline in 2016 from Glastir Monitoring and Evaluation Program data under predicted climate data (UKCP18).

**Figure SM.8.**
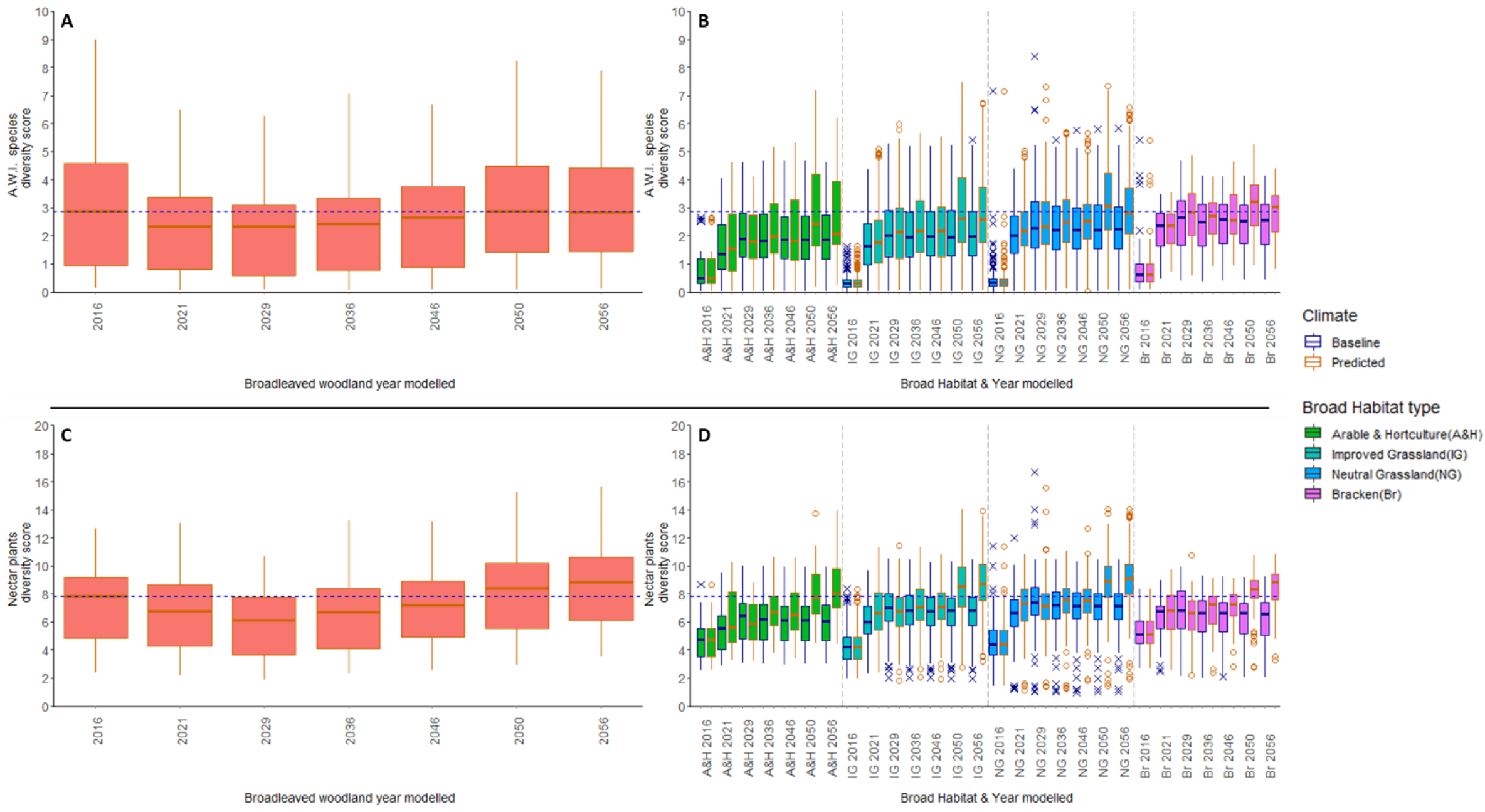
Boxplots of species group diversity scores: A, Broadleaved woodland habitat ancient woodland indicator (A.W.I.) species at baseline in 2016 and later years with predicted climate; B, the four habitats A.W.I. species scores at baseline and modelled as being planted broadleaf in subsequent years, with baseline and predicted climate; D Broadleaved woodland habitat nectar producing (N.P.) species at baseline in 2016 and later years with predicted climate; C the four habitats N.P. species scores at baseline and modelled as being planted broadleaf in subsequent years, with baseline and predicted climate. The dashed blue line represents the median A.W.I. score for broadleaved woodland at baseline in A&B and the same N.P. mean for D&C. The data used to create these boxplots was generated using an ecological niche model MultiMOVE, inputs were altered to represent baseline (1981-2016) and future climates using downscale UKCP18 climate data, incremental increase of cover weighted canopy height representing tree growth and generalised linear mixed effect models of soil variable change under broadleaved plantation.

**Figure SM.9.**
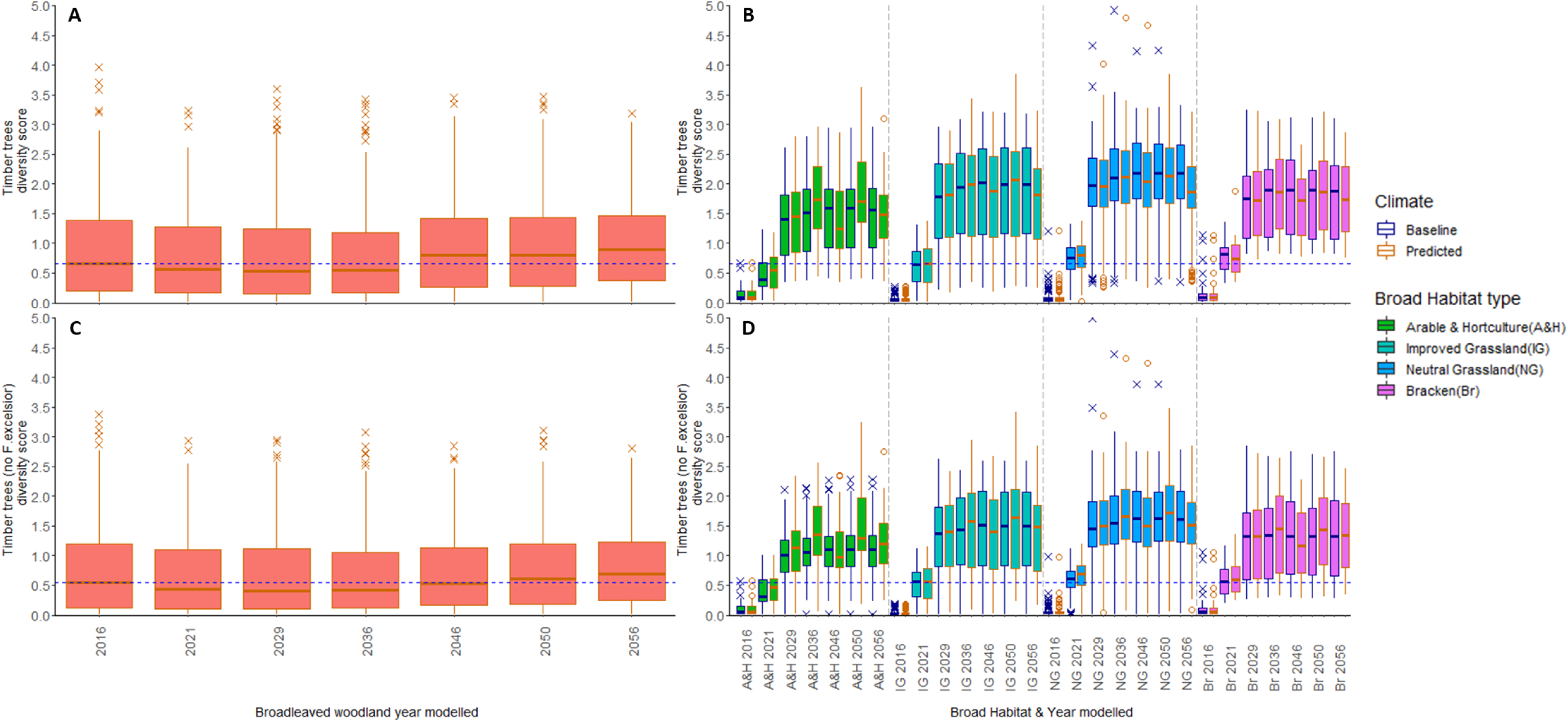
Boxplots of diversity scores of timber species group diversity (TSGD): A, Broadleaved woodland habitat TSGD at baseline in 2016 and later years with predicted climate; B, the four habitats at baseline and modelled as being planted broadleaf TSGD in subsequent years, with baseline and predicted climate; D is the same as A but with *Fraxinus excelsior* removed; C is the same as B with *Fraxinus excelsior* removed. The dashed blue line represents the mean TSGD for broadleaved woodland at baseline in A&B and the mean TSGD without *F. excelsior* for broadleaved woodland at baseline for D&C. The data used to create these boxplots was generated using an ecological niche model MultiMOVE, inputs were altered to represent baseline (1981-2016) and future climates using downscale UKCP18 climate data, incremental increase of cover weighted canopy height representing tree growth and generalised linear mixed effect models of soil variable change under broadleaved plantation.

**Figure SM.10.**
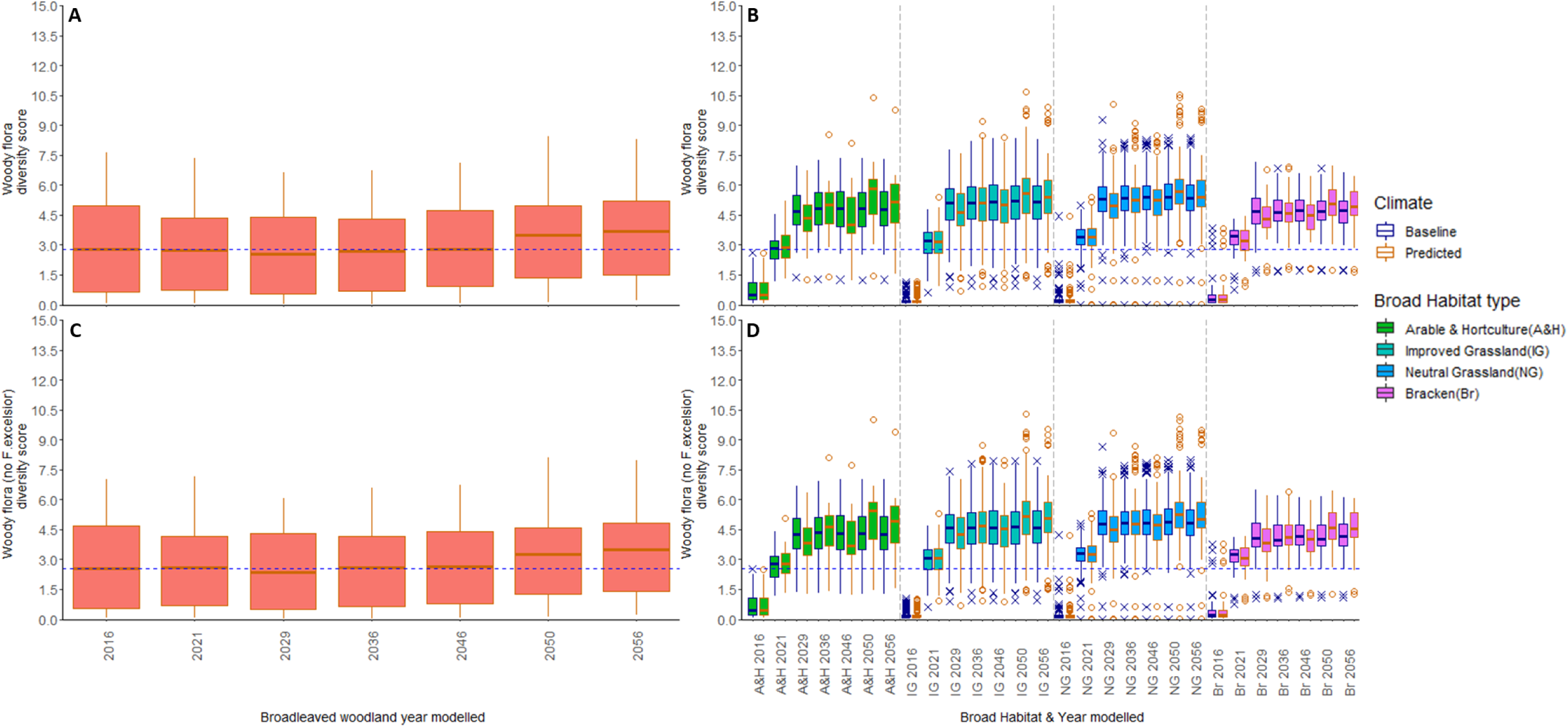
Boxplots of diversity scores of woody flora (WFD): A, Broadleaved woodland habitat WFD at baseline in 2016 and later years with predicted climate; B, the four habitats at baseline and modelled as being planted broadleaf WFD in subsequent years, with baseline and predicted climate; D is the same as A but with *Fraxinus excelsior* removed; C is the same as B with *Fraxinus excelsior* removed. The dashed blue line represents the median WFD for broadleaved woodland at baseline in A&B and the median WFD without *F. excelsior* for broadleaved woodland at baseline for D&C. The data used to create these boxplots was generated using an ecological niche model MultiMOVE, inputs were altered to represent baseline (1981-2016) and future climates using downscale UKCP18 climate data, incremental increase of cover weighted canopy height representing tree growth and generalised linear mixed effect models of soil variable change under broadleaved plantation.

**Figure SM.11.**
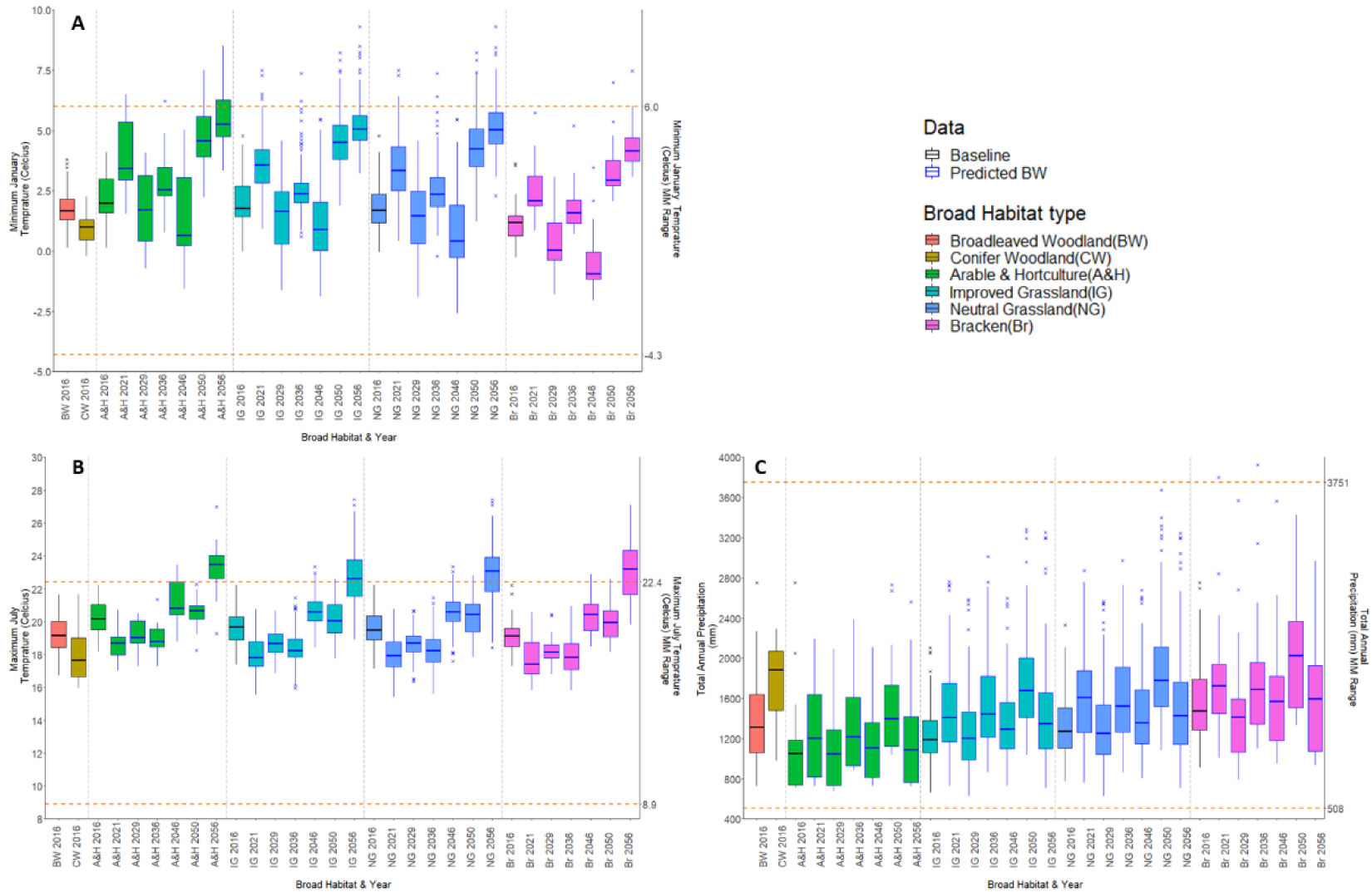
Boxplots showing Glastir Environmental Monitoring Program X-plot climate data from UKCP18 at 1 km square scale from 2016 (averaged from 1981 to 2016 as a baseline) through subsequent predicted years up to 2056. Observed (2016, black edged box plots) data was sourced from Met Office HadUK-Grid, 1 km climate data and averaged. Predictions data (blue edged box plots) from the UKCP18 high emissions scenario, RCP8.5, UK regional 12 km scale probabilistic data, was downscaled to 1km. A. Total annual precipitation in mm, B. minimum January temperature in °C and C. Maximum July temperature in °C; georeferenced to the 1 km square each X-plot was within. Left hand Y-axes shows variable ranges to derive boxplot values from (black observed average for 2016; blue beyond 2016 predicted). The dashed orange lines show the top and bottom of ranges the R package MultiMOVE was constructed within. X-axis labels show the X-plot groups of year and board habitat type: A&H = arable and horticulture; I.G. = improved grassland; N.G. = neutral grassland; Br = Bracken.

## SM 4 | Climatic variables

### SM 4.1 | Downscaled climatic variables

The 1 km climate variables for future projections were taken from a single member (01) of the CHESS-SCAPE ensemble (Robinson *et al*., 2022). This was downscaled from the corresponding member (01) of the UKCP18 regional climate model perturbed parameter ensemble (RCM-PPE) (Met Office Hadley Centre, 2018). This is an ensemble of RCM variants, nested within perturbed parameter variants of the HadGEM3-GC3.05 global climate model (GCM) (Murphy *et al.,* 2018). This nesting allows better projection of the dynamics of regional UK and European climate without the prohibitive computational cost of running the model globally at the high resolution. Ensemble member 01 uses the default model parameters (Murphy *et al.,* 2018) and CO^2^ concentrations prescribed by RCP8.5 (van Vuuren *et al*., 2011). The RCM-PPE data are distributed at 12 km resolution. To produce CHESS-SCAPE, these were then downscaled to 1 km using an adapted version of the CHESS methodology (Robinson *et al*., 2017), which interpolates variables to a finer resolution while adjusting for local topography using physically-based and empirical methods (Robinson *et al*., in prep.). The resulting files cover the UK land surface, but exclude Shetland due to data availability. Variables were reduced from the grid box elevation of the climate model to mean sea level, interpolated from 12 km to 1 km, then readjusted to the elevation at 1 km resolution given by the Integrated Hydrological Digital Terrain Model (IHDTM) (Morris *et al*., 1990). Daily mean, minimum and maximum air temperatures were adjusted using a lapse rate of -0.006 K m^-1^ (Hough & Jones, 1997), and the interpolation was carried out separately for each variable. Rainfall was not interpolated but was adjusted for long-term averages in local rainfall rates using a 1 km resolution map of Standardised Area Average Rainfall (Spackman, 1993). In this study we use the climate projection without bias-correction.

The RCM-PPE has a strong climate sensitivity and is at the high end of the range of the CMIP5 ensemble, but it is consistent with the current generation of climate models in CMIP6 (Lowe *et al*., 2018), particularly the related model HadGEM3-GC.3.1 (Williams *et al*., 2017). Although the overall trend is for warming temperatures, the interannual variability of the climate model projection is such that 2029 is cooler than 2026. Additionally, there was a particularly warm period in the climate model projection from 2017 to 2027, which resulted in 2026 being outside of the baseline range. While this is unusual, it is consistent with climate variability. The projected trend in rainfall is for an overall decrease in mean annual rainfall by 2080 (with an increase in winter rainfall but a bigger decrease in summer rainfall), although there is little change by the end of the study period (Murphy *et al.,* 2018). Again, interannual variability can be seen with higher rainfall in 2021 and 2026, followed by lower in 2029.

### SM 4.2 | Modelling into future climate space unknowns

Given the substantial unknowns of attempting this type of work into the future under global change the ecological modelling community has a challenge to achieve reliable predictions regardless of the methods used (Williams *et al*., 2017; Williams *et al*., 2007; Veloz *et al*., 2012; Smith *et al*., 2013; Fitzpatrick & Hargrove, 2009). Considering novel environmental space occurrence into the future, we do not attempt to predict climate effects on biogeochemical processes here e.g. carbon priming effects (Smith *et al*., 2013) as we restrain consideration of results to those that do not fall outside MultiMOVE operating space thus is not expected to be impactful. A greater concern maybe that early successional or disturbed ecosystems are at greater risk to climate change impacts (Kröel-Dulay *et al*., 2015) this maybe be mitigated by consideration of species beyond natives but that are likely to be ecological functional and non-damaging e.g. tree species examples in Read *et al*. (2009).

## SM 5 | Further conclusive text

### SM 5.1 | F. excelsior removal from the species pool

The removal of *F. excelsior* from the species pool is a simplification of reality as the trees would be more likely to decline over time with *H. fraxineus* infection (Pautasso *et al*., 2013; Mitchell *et al*., 2016; Skovsgaard *et al*., 2017). However, we have not applied an incremented reduction in *F. excelsior*. This is due to no data being found on *H. fraxineus* effects on *F. excelsior* abundance with sufficient robustness to estimate change, despite our searching. This is an area that may merit further research to measure dieback impacts at largescale and long-term. Therefore, removal of *F. excelsior* from the plantation scenarios seems to be an at least adequate representation of possible future realities.

### SM 5.2 | Management and Policy

To pursue forest establishment for biodiversity loss mitigation and net zero 2050 goals with climate change and tree disease being influential factors, we suggest the following points for management and policy decisions. The 30-50 year time-frame for successional forest establishment that abandonment / no intervention rewilding takes, in comparison with the 20-40 years taken for some variables modelled to reach broadleaved woodland baseline values, does suggest that in the right context for certain outcomes planting can be a more rapid method of afforestation. To give UK forest ecosystems the best opportunities to establish we make 5 recommendations for management and policy:

1. Species selection in its simplest form can be done by local observation (10 km) of desirable endpoint forest habitats via tree species matching, but would likely be best done by consideration of site environmental conditions or species selection tools e.g. timber species suggested via the Forestry Commission’s ESC tool, URL: http://www.forestdss.org.uk/geoforestdss/ (Pyatt, Ray and Fletcher, 2001).
2. Planting method must be of minimum soil disturbance to reduce and possible carbon loss e.g. by spade and hand as suggested in Berdeni *et al*. (2021), this could also reduce negative carbon priming effects (De Graff *et al*., 2006) especially if this practice happens at scale.
3. Planting adjacent to or in close proximity to established woodland (especially ancient or long term) as older sites are far more likely to have desirable species that will colonise as plantations establish to forests (Brunet, De Frenne, Holmström, & Mayr, 2012; Di Sacco & Hardwick *et al*., 2020; Thomaes *et al*., 2012).
4. Consideration of “most creation” versus “most change” for example planting up a neutral grassland site adjacent to an ancient woodland may provide the “most creation” but greater biodiversity and soil condition recovery (“most change”) may come from planting an adjacent arable field.
5. Lastly policy and legislative support (including forest planning support) as it is unreasonable to expect land-managers to hold the necessary ecological knowledge to make decisions that are, economically viable for them as well as mitigating biodiversity loss and climate change. While policy documentation to some extend does acknowledge this (Davies, 2016; Defra, 2018), legislation or supportive schemes directly applied to ecosystem management (rather than land management) is rare. However, the new UK Environmental Land Stewardship scheme (ELMs) presents an opportunity to change this (Defra, 2020).

