## Supplementary material for "Make like a tree and leave: How will tree species loss and climate change alter future temperate broadleaved forests?": Neural_net_building_files_V2.0: Building_the_neural_nets_notebook.Rmd

````
---
title: "Creating neural network models that translate soil variables into mean Ellenberg values"
author: "S_Smart"
date: "23/06/2021"
output: html_notebook
---

```{r setup, include=FALSE}
knitr::opts_chunk$set(echo = TRUE)

## setting working directory.. change this if new user
knitr::opts_knit$set(root.dir = "C:\\Bedes_paper_2021")

```

# Aim

Here we build models to produce estimates of mean Ellenberg values for soil moisture, pH and fertility given soil %C, %N and gravimetric moisture. A re-working could usefully add in canopy height too as we know shade is a also an influential filter on the way the species assemblage responds to other abiotic gradients.

## Building the Neural Nets

The code below derives a new calibration of mean Ellenbergs given soil data. The resulting neural network models can then be applied in predictive mode given new soil inputs. To build the models we use the [Countryside Survey](https://countrysidesurvey.org.uk/) 2000 dataset of mean Ellenberg scores based on plant species composition (not weighted by cover/abundance) and co-located soil data that were used to produce the first set of transfer functions reported in [Smart et al (2010)](https://onlinelibrary.wiley.com/doi/abs/10.1111/j.1654-1103.2010.01173.x).

We train models on a random 70% and test on a random 30% of the data. First we need to carry out a sensitivity analysis to determine the best number of hidden layers to use in our black-box models. We selected neural nets because we have less interest in the form of the regression relationships between predictors and response but more interest in generating a model that can best fit training and new independent test data. The last step is critical because we need to ensure transferability to new areas and samples.

Some of the following code was modified from: <http://www.michaeljgrogan.com/neural-network-modelling-neuralnet-r/>

## How many hidden layers and neurons do we need?

"There are some empirically-derived rules-of-thumb, of these the most commonly relied on is 'the optimal size of the hidden layer is usually between the size of the input and size of the output layers'. Jeff Heaton, author of 'Introduction to Neural Networks in Java' offers a few more. In sum, for most problems, one could probably get decent performance (even without a second optimization step) by setting the hidden layer configuration using just two rules: (i) number of hidden layers equals one; and (ii) the number of neurons in that layer is the mean of the neurons in the input and output layers."

So here, we experiment with 1 or 2 hidden layers and with 2 or 3 neurons in the first layer. See here for further information

<https://stats.stackexchange.com/questions/181/how-to-choose-the-number-of-hidden-layers-and-nodes-in-a-feedforward-neural-network>

```{r load packages}
library(neuralnet) # NOTE THAT 'COMPUTE' HAS BEEN REPLACED WITH 'PREDICT'...PFFFFF!
library(caTools)
library(dplyr)
library(haven)
library(ggplot2)

```

Loop to select best Neural network model based on different number of neurons and hidden layers and using a number of measures of model performance.

The search for the best model is done separately for each mean Ellenberg score.

## Build the best models

```{r model building}

# Load the CS1998 data used to produce the original transfer functions.

Soils_and_Ebergs98<- read.csv("CS98_input.csv")

Input<-Soils_and_Ebergs98[,c(3:5,8,12)]

# Create dataframe to take testing stats
   
   L1 <- (1:4) # Here we test combinations of 1 to 2 hidden layers each with different numbers of neurons and select the best perfroming model
   L2 <- (0:2)
   d1 <- expand.grid(L1 = L1, L2 = L2)
   
   Net_results<-data.frame(d1, accuracy=rep(0,12), mEb_diff=rep(0,12),
                           Net_code=(1:12))

   # Create normalization function  
   
   normalize <- function(x) {
     return ((x - min(x)) / (max(x) - min(x)))
   }

      #MAX-MIN NORMALIZATION
   
   maxmindf <- as.data.frame(lapply(Input, normalize))
   
   #TRAINING AND TEST DATA - just uses caTools 
   
   set.seed(101)
   
   # Create Split (any column is fine)
   split = sample.split(maxmindf$EbW, SplitRatio = 0.70)
   
   # Split based off of split Boolean Vector
   train = subset(maxmindf, split == TRUE)
   test = subset(maxmindf, split == FALSE)
   
   #4. NEURAL NETWORK
   #4.1 Build formula 
   
   var_names <- names(train)
   
   # Concatenate strings
   f <- paste(var_names[c(1:4)], collapse=' + ')
   f <- paste('EbW ~',f)
   
   # Convert to formula
   f <- as.formula(f)
   
   # Experiment with hidden layers
   
   for(i in (Net_results$Net_code)){
     
          h1=Net_results$L1[i]
     h2=Net_results$L2[i]
     
     if (h2==0) {nn <- neuralnet(f,train, hidden=c(h1), linear.output=TRUE,stepmax=1e6)
     } else {nn <- neuralnet(f,train, hidden=c(h1,h2), linear.output=TRUE,stepmax=1e6)}
    
   # Model Validation
   
   # Compute Predictions off Test Set
   predicted.nn.values <- compute(nn, test[1:4])
   
   results <- data.frame(actual = test$EbN, prediction = predicted.nn.values$net.result)
  
   # Generate absolute deviation of predicted versus observed
   predicted=results$prediction * abs(diff(range(Input$EbN))) + min(Input$EbN)
   actual=results$actual * abs(diff(range(Input$EbN))) + min(Input$EbN)
   comparison=data.frame(predicted,actual)
   deviation=((actual-predicted)/actual)
   abs_deviation=(actual-predicted)
   comparison=data.frame(predicted,actual,deviation,abs_deviation)
   Net_results$mEb_diff[i]<-mean(abs(abs_deviation))
   Net_results$accuracy[i]<-1-abs(mean(deviation))
   
   }

   plot(comparison$predicted, comparison$actual) 
   
# Results table exported.. best model has just one hidden layer with one neuron!
   write.csv(Net_results, file = "EbN_nnet_tests.csv")
   
   
   # Reproduce best model
   nn <- neuralnet(f,train, hidden=c(2), linear.output=TRUE,stepmax=1e6)
   
   
   # save best model object ready to use for prediction
   save(EbW, file = "nn_EbW.rda")

```

# Independent testing against data from the Welsh GMEP survey carried out between 2013-'16

Here we compare the predictive performance of the original GBMOVE equations in [Smart et al (2010)](https://onlinelibrary.wiley.com/doi/abs/10.1111/j.1654-1103.2010.01173.x) with the hopefully better performance of the neural nets.

We do each Ellenberg score in sequence as follows:

## 1. Ellenberg Wetness scores:

```{r test Ellenberg wetness}

#load NNet models
   load(file = "nn_EbW.rda")
   
   #Read GMEP data for model testing  
   GMEP<- read.csv(Test_gmep_soils_bergsx1.csv")
   Test<-GMEP[,c(4,13,10,9,11)]
   
   # Calculate Ebergs based on GBMOVE calibration formulae
   
   Test$GbmW <- (log((Test$MC/(100-Test$MC)))+3.27)/0.55
   
   # Examine
   plot(Test$GbmW, Test$EbW)
   # Delete NAs and preds outside range
   
   Test1<-subset(Test, GbmW<=12 & GbmW>=1)
   plot(Test1$EbW, Test1$GbmW)
   
   # Now solve using NNet
   
   #MAX-MIN NORMALIZATION
   normalize <- function(x) {
     return ((x - min(x)) / (max(x) - min(x)))
   }
   normTest <- as.data.frame(lapply(Test1, normalize))
   
  
   # Compute Predictions off Test Set
   predicted.EbW.values <- compute(EbW, normTest[2:5])
   
   results <- data.frame(obs = Test1$EbW, GLM_EbW=Test1$GbmW, prediction = predicted.EbW.values$net.result)
   # Note here that the back-transformation requires the values of range and min from the 
   # original dataset used to create the neural net - see below for a list of these.
    
   results$predicted<-(results$prediction * 4.866667) + 4.333333
   
   # Now calculate the separate diagnistic stats. Needed for each comparison of 
   # obs v GLM_Eb* and obs v predicted
   
   results$GLM_deviation<-((results$obs-results$GLM_EbW)/results$obs)
   results$GLM_abs_deviation<-(results$obs-results$GLM_EbW)
   
   GLMmEb_diff=mean(results$GLM_abs_deviation)
   GLMaccuracy=1-abs(mean(results$GLM_deviation))
   
   results$NN_deviation<-((results$obs-results$predicted)/results$obs)
   results$NN_abs_deviation<-(results$obs-results$predicted)
   
   NNmEb_diff=mean(results$NN_abs_deviation)
   NNaccuracy=1-abs(mean(results$NN_deviation))
  
   GLMaccuracy
   GLMmEb_diff
   NNaccuracy
   NNmEb_diff
   
plot(results$obs, results$predicted) 
plot(results$obs, results$GLM_EbW)  


```

```{r model building and selection}


```

|                                                 | Neural Nets | GLM (Smart et al 2010) |
|-------------------------------------------------|-------------|------------------------|
| \% agreement pred v observed                    | 0.96        | 0.86                   |
| Mean deviation in Ellenberg scores (obs v pred) | 0.30        | 0.79                   |

Higher accuracy with the Neural net and much lower average absolute difference in mean Ellenberg values such that the difference between obersved and predicted is on average 0.3 of an Ellenberg unit.

## 2. Ellenberg N scores:

```{r Ellenberg N}


#load NNet models
load(file = "nn_EbN.rda")

#Read GMEP data for model testing  
GMEP<- read.csv("Test_gmep_soils_bergsx1.csv")
Test<-GMEP[,c(2,13,10,9,11)]

# Calculate Ebergs based on GBMOVE calibration formulae

Test$GbmN <- exp(0.7751 - (0.00006*Test$MC) - (0.00009*(Test$MC^2)) - (0.01475*Test$C) + (0.000099*(Test$C^2)) + (0.2639*Test$pH) 
                    - (0.01684*(Test$pH^2)) + (0.1908*Test$N))

# Examine
plot(Test$GbmN, Test$EbN)

# Now solve using NNet

#MAX-MIN NORMALIZATION
normalize <- function(x) {
  return ((x - min(x)) / (max(x) - min(x)))
}
normTest <- as.data.frame(lapply(Test, normalize))


# Compute Predictions off Test Set
predicted.EbN.values <- compute(EbN, normTest[2:5])

results <- data.frame(obs = Test$EbN, GLM_EbN=Test$GbmN, prediction = predicted.EbN.values$net.result)
# Note here that the back-transformation requires the values of range and min from the 
# original dataset used to create the neural net - see below for a list of these.

results$predicted<-(results$prediction * 6.083333) + 1.166667

# Now calculate the separate diagnistic stats. Needed for each comparison of 
# obs v GLM_Eb* and obs v predicted

results$GLM_deviation<-((results$obs-results$GLM_EbN)/results$obs)
results$GLM_abs_deviation<-(results$obs-results$GLM_EbN)

GLMmEb_diff=mean(results$GLM_abs_deviation)
GLMaccuracy=1-abs(mean(results$GLM_deviation))

results$NN_deviation<-((results$obs-results$predicted)/results$obs)
results$NN_abs_deviation<-(results$obs-results$predicted)

NNmEb_diff=mean(results$NN_abs_deviation)
NNaccuracy=1-abs(mean(results$NN_deviation))

GLMaccuracy
GLMmEb_diff
NNaccuracy
NNmEb_diff


plot(results$obs, results$predicted) 
plot(results$obs, results$GLM_EbN)

```

|                                                 | Neural Nets | GLM (Smart et al 2010) |
|-------------------------------------------------|-------------|------------------------|
| \% agreement pred v observed                    | 0.88        | 0.85                   |
| Mean deviation in Ellenberg scores (obs v pred) | -0.25       | -0.29                  |

Neural network model outperforms the GLM on both counts but there is less to separate them than for the Ellenberg W models above.

## 3. Ellenberg R (pH) scores

```{r Ellenberg R}


#load NNet models load(file = "nn_EbR.rda")

#Read GMEP data for model testing\
GMEP<- read.csv("Test_gmep_soils_bergsx1.csv") 
Test<-GMEP[,c(3,13,10,9,11)]

# Calculate Ebergs based on GBMOVE calibration formulae


Test$GbmR <- 0.5293 - (0.02503*Test$MC) + (1.665*Test$pH) - (0.1061(Test$pH^2)) - (0.00566*Test$C) 


# Examine

plot(Test$GbmR, Test$EbR)

# Now solve using NNet

#MAX-MIN NORMALIZATION normalize \<- function(x) { return ((x - min(x)) / (max(x) - min(x))) } normTest \<- as.data.frame(lapply(Test, normalize))

# Compute Predictions off Test Set

predicted.EbR.values <- predict(EbR, normTest[2:5])

results <- data.frame(obs = Test$EbR, GLM_EbR=Test$GbmR, prediction = predicted.EbR.values$net.result) 

# Note here that the back-transformation requires the values of range and min from the  original dataset used to create the neural net - see below for a list of these.

results$predicted<-(results$prediction * 5.25) + 2

# Now calculate the separate diagnistic stats. Needed for each comparison of

# obs v GLM_Eb\* and obs v predicted

results$GLM_deviation<-((results$obs-results$GLM_EbR)/results$obs) results$GLM_abs_deviation<-(results$obs-results$GLM_EbR)

GLMmEb_diff=mean(results$GLM_abs_deviation) GLMaccuracy=1-abs(mean(results$GLM_deviation))

results$NN_deviation<-((results$obs-results$predicted)/results$obs) results$NN_abs_deviation<-(results$obs-results$predicted)

NNmEb_diff=mean(results$NN_abs_deviation) NNaccuracy=1-abs(mean(results$NN_deviation))

GLMaccuracy 
GLMmEb_diff 
NNaccuracy 
NNmEb_diff


plot(results$obs, results$predicted) 
plot(results$obs, results$GLM_EbR)


```

|                                                 | Neural Nets | GLM (Smart et al 2010) |
|-------------------------------------------------|-------------|------------------------|
| \% agreement pred v observed                    | 0.90        | 0.84                   |
| Mean deviation in Ellenberg scores (obs v pred) | -0.28       | -0.41                  |

The range of the GLM predictions is substantially narrower than observed values although accuracy does not differ much hugely compared to the neural net largely because of the residual variation around the observed values between EbR = 3 and 5. However, on balance the neural net is again better. The neural net predictions also have a much lower average absolute difference from the observations.
````
